# OTULIN prevents liver inflammation and hepatocellular carcinoma by inhibiting FADD- and RIPK1 kinase-mediated hepatocyte apoptosis

**DOI:** 10.1101/776088

**Authors:** Lien Verboom, Arne Martens, Dario Priem, Esther Hoste, Mozes Sze, Hanna Vikkula, Sofie Voet, Laura Bongiovanni, Alain de Bruin, Charlotte L. Scott, Manolis Pasparakis, Mathieu JM Bertrand, Geert van Loo

## Abstract

Inflammatory signaling pathways are tightly regulated to avoid chronic inflammation and the development of inflammatory pathologies. OTULIN is a deubiquitinating enzyme that specifically cleaves linear ubiquitin chains generated by the linear ubiquitin chain assembly complex (LUBAC), and OTULIN deficiency causes OTULIN-related autoinflammatory syndrome (ORAS) in humans. OTULIN was shown to negatively control NF-κB signaling in response to various stimuli, but also to protect cells from tumor necrosis factor (TNF)-induced apoptosis. To investigate the importance of OTULIN in liver homeostasis and pathology, we developed a novel mouse line specifically lacking OTULIN in liver parenchymal cells. These mice spontaneously develop a severe liver disease, characterized by liver inflammation, hepatocyte apoptosis and compensatory hepatocyte proliferation, leading to steatohepatitis, fibrosis and hepatocellular carcinoma (HCC). Genetic ablation of Fas-associated death domain (FADD) completely rescues the severe liver pathology, and knock-in expression of kinase inactive receptor-interacting protein kinase 1 (RIPK1) significantly protects from developing liver disease, demonstrating that death receptor-mediated apoptosis of OTULIN-deficient hepatocytes triggers disease pathogenesis in this model. Finally, we demonstrate that type I interferons contribute to disease pathogenesis in hepatocyte-specific OTULIN deficient mice. Together, our study reveals the critical importance of OTULIN in protecting hepatocytes from death, and thereby avoid development of chronic liver inflammation and HCC in mice.

## Introduction

Liver cancer is the second most frequent cause of cancer-related deaths and the fifth most common type of cancer world-wide (Ringelhan et al., 2018). From all primary liver cancers, hepatocellular carcinoma (HCC) is the most frequent, responsible for 80-90 % of all cases, and nearly all develop as a result of chronic liver inflammation inducing fibrosis, cirrhosis and finally HCC. There is clear evidence that cell death represents a basic biological process in liver cancer, where hepatocyte death induces compensatory hepatocyte regeneration, chronic liver inflammation and activation of non-parenchymal cells, promoting liver fibrosis and tumorigenesis (Luedde and Schwabe, 2011; Kondylis and Pasparakis, 2019).

Tumor necrosis factor (TNF) is a major inflammatory cytokine, capable of inducing inflammatory gene expression, but also of triggering cell death. Many studies have shown that optimal regulation of TNF signaling is essential to maintain liver homeostasis and prevent liver inflammation and inflammation-induced HCC (Luedde et al., 2014). Inflammatory signaling, such as by TNF, is heavily controlled by ubiquitination, a posttranslational modification of proteins (Iwai et al., 2014). Sensing of TNF by TNF receptor 1 (TNFR1) initiates the assembly of a receptor-proximal complex, known as complex I, which activates the mitogen activated protein kinase (MAPK) and nuclear factor-κB (NF-κB) signaling pathways (Ting and Bertrand, 2016). The initial binding of TNFR1-associated death domain protein (TRADD) and receptor-interacting protein kinase 1 (RIPK1) to the receptor allows the subsequent recruitment of TNF receptor-associated factor 2 (TRAF2) and of the E3 ubiquitin ligases cellular inhibitors of apoptosis 1 and 2 (cIAP1/2). The K63-ubiquitin chains generated by cIAP1/2 serve as docking stations for the adaptor proteins TAB2/3 and for the recruitment of the kinase TAK1, which subsequently activate the MAPK signaling pathways. These K63-ubiquitin chains also help recruiting the multiprotein E3 ubiquitin ligase complex LUBAC (composed of HOIL-1, HOIP and SHARPIN), which further conjugates complex I components with linear-ubiquitin chains. The adaptor protein NEMO binds to these linear chains and brings the kinases IKKα and IKKβ to the complex, thereby allowing activation of the NF-κB pathway. NEMO recruitment to the complex also enables activation of the kinases TBK1/IKKε, which may regulate NF-κB activation (Clark et al., 2011; Lafont et al., 2018; Xu et al., 2018). Upon activation, the MAPK and NF-κB pathways drive expression of a large set of genes, including pro-inflammatory genes (Ting and Bertrand, 2016).

The linear-ubiquitin chains generated by LUBAC in the TNFR1 pathway are therefore essential for the NF-κB-dependent expression of pro-inflammatory genes, but were also shown to play a double and crucial role in preventing TNF cytotoxicity. Indeed, apart from pro-inflammatory molecules, activation of NF-κB also leads to the transcriptional upregulation of anti-apoptotic proteins, which protect cells from RIPK1 kinase-independent apoptosis (Wang et al., 2008). In addition, the linear ubiquitin-dependent phosphorylation of RIPK1 by IKKα/β- and TBK1/IKKε was shown to protect cells from RIPK1 kinase-dependent apoptosis (Dondelinger et al., 2015, 2019; Lafont et al., 2018; Xu et al., 2018; Ting and Bertrand, 2016). Consequently, when these protective brakes are compromised, such as following LUBAC deficiency (Peltzer et al., 2014, 2018) (Priem et al. CDDis 2019), activated TRADD and RIPK1 dissociate from complex I, and respectively induce RIPK1 kinase-independent and -dependent apoptotic cascades through association with Fas-associated death domain (FADD) protein and procaspase-8 to form the cytosolic death inducing complex II. Necroptosis occurs when caspase-8 activation is blocked, and involves RIPK1 kinase-dependent recruitment of RIPK3 and mixed lineage kinase domain-like protein (MLKL) to complex II, inducing strong inflammatory responses (Ting and Bertrand, 2016). Hence, LUBAC activity is essential to prevent apoptotic and necroptotic cell death, and HOIP and HOIL-1 deficient mice die during development (Peltzer et al., 2014, 2018).

Ubiquitination is a reversible process, and the deubiquitinating (DUB) enzyme OTULIN (OTU deubiquitinase with linear linkage specificity, also known as Fam105b or Gumby) exclusively cleaves the linear ubiquitin chains generated by LUBAC, and hence critically controls inflammatory and cell death responses (Keusekotten et al., 2013; Rivkin et al., 2013). Accordingly, mice harboring a point mutation abolishing the ability of OTULIN to bind ubiquitin (so-called ‘gumby’ mice) or knockin mice expressing a catalytically inactive variant of OTULIN die in utero due to excessive inflammatory cell death (Rivkin et al., 2013; Heger et al., 2018). However, mice with inducible OTULIN deficiency were shown to be viable and develop a severe inflammatory disease (Damgaard et al., 2016), resembling an autoinflammatory syndrome in humans carrying homozygous hypomorphic OTULIN mutations, called OTULIN-related autoinflammatory syndrome (ORAS, also known as Otulipenia) (Zhou et al., 2016; Damgaard et al., 2016, 2019; Nabavi et al., 2019). Although OTULIN was originally shown to negatively regulate LUBAC activity, its deficiency does not induce LUBAC hyperactivity, as expected, but rather suppresses its function. Indeed, OTULIN, HOIP and HOIL-1 deficient mice have very similar phenotypes displaying embryonic lethality due to uncontrolled cell death. However, OTULIN turned out to promote rather than counteract LUBAC signaling by preventing LUBAC autoubiquitination through the removal of linear ubiquitin chains from LUBAC (Heger et al., 2018).

In light of the reported function of OTULIN in regulating both TNF-induced NF-κB signaling and cell death (Keusekotten et al., 2013; Damgaard et al., 2016, 2019; Heger et al., 2018), we questioned the role of OTULIN in liver (patho)physiology. For this, we generated OTULIN conditional knockout mice that are specifically deficient for OTULIN in liver parenchymal cells. These mice spontaneously develop chronic liver inflammation, fibrosis and hepatocellular carcinomas, illustrating the importance of OTULIN in normal liver tissue homeostasis. Chronic liver pathology results from the hypersensitivity of OTULIN-deficient hepatocytes to spontaneous FADD- and RIPK1 kinase activity-dependent apoptosis, which triggers compensatory hepatocyte proliferation and inflammation. Interestingly, type I interferon (IFN) signaling also contributes to liver pathology in OTULIN deficient mice. Together, these studies establish OTULIN as a crucial hepatoprotective factor.

## Results

### Development of a severe liver pathology in hepatocyte-specific OTULIN knockout mice

*Otulin*-targeted ES cells (Otulin^tm1a(EUCOMM)Hmgu^) were used to generate chimeric mice that transmitted the targeted allele to their offspring. Mice homozygous for the LoxP-flanked *Otulin* allele (*Otulin*^FL/FL^) express normal levels of OTULIN and develop normally (data not shown). Deletion of the LoxP-flanked *Otulin* alleles through expression of a ubiquitously expressed Cre recombinase leads to a loss of OTULIN protein as shown in mouse embryonic fibroblasts (MEF) (Suppl. Fig. 1A). In agreement with previous studies (Rivkin et al., 2013; Damgaard et al., 2016; Heger et al., 2018), full body OTULIN knockout mice are not viable (Suppl. Fig. 1B), preventing further studies. In order to study the role of OTULIN in hepatocytes and in liver physiology and pathology, we crossed the *Otulin*^FL/FL^ mice with a transgenic mouse line that expresses Cre under the control of the liver-specific albumin/alpha-fetoprotein promoter/enhancer (Alfp-Cre) which mediates efficient Cre recombination in liver parenchymal cells (Kellendonk et al., 2000) (Suppl. Fig. 1C-D). Hepatocyte-specific OTULIN knockout (Otulin^FL/FL^/Alfp-Cre, liver parenchymal cell-specific OTULIN knockout, OTULIN^LPC-KO^) mice were born with normal Mendelian segregation. Immunoblot analysis of liver protein extracts revealed efficient ablation of OTULIN in livers of OTULIN^LPC-KO^ mice (Fig. 1A). Residual OTULIN expression in these livers can be attributed to nonparenchymal liver cells that are not targeted by the Alfp-Cre allele.

**Figure 1.**
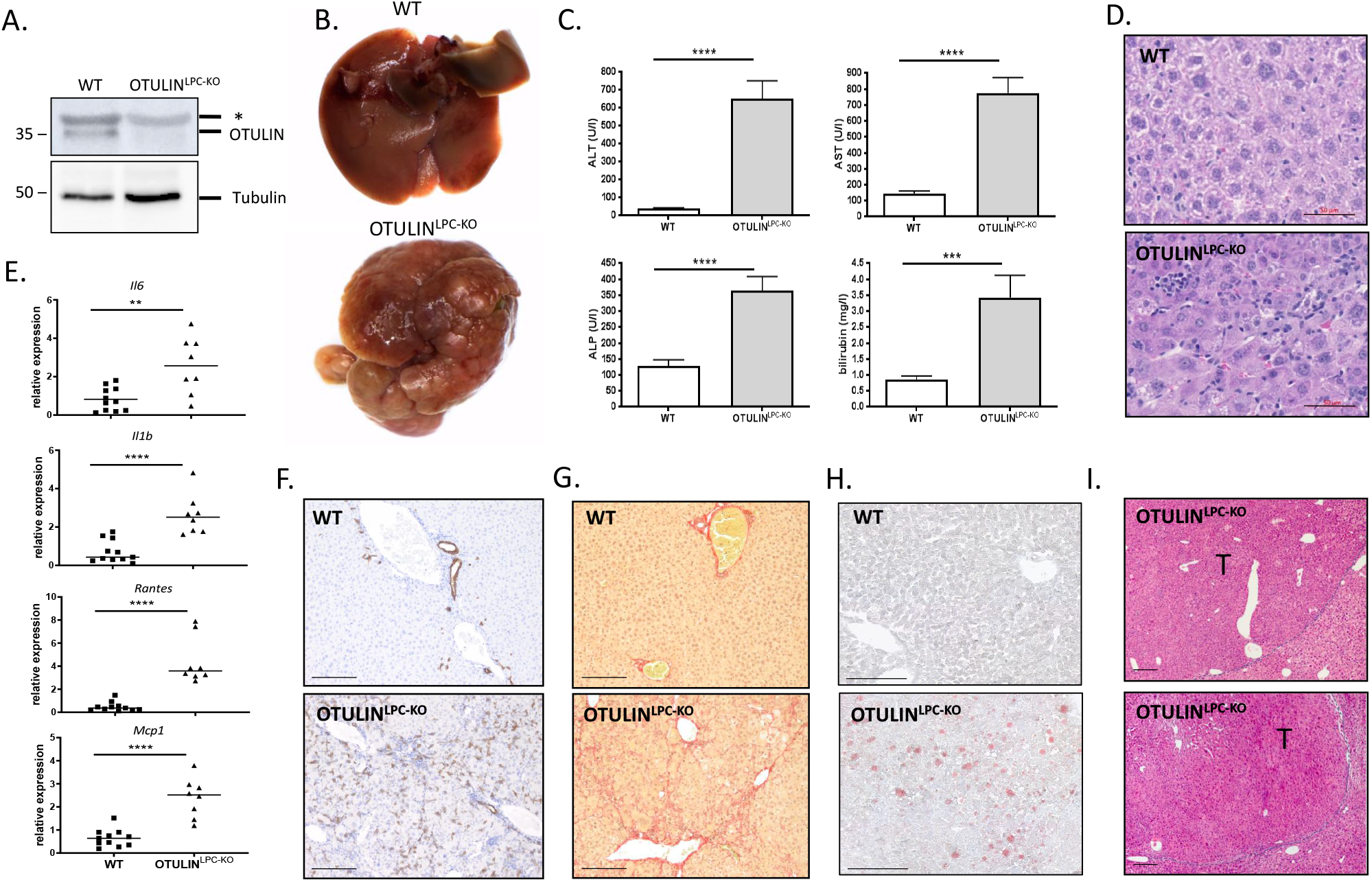
Development of a severe liver phenotype in OTULIN^LPC-KO^ mice. (A) Western blot analysis for OTULIN expression in total liver lysates from a control wild-type (WT) and OTULIN^LPC-KO^ littermate mouse. Anti-tubulin immunoblotting was used as loading control. *, unspecific. Data are representative of three independent experiments. (B) Macroscopic pictures of representative livers from a 10 week-old wild-type (WT) and OTULIN^LPC-KO^ littermate mouse. (C) Serum ALT, AST, ALP and bilirubin levels of control (n = 11) and OTULIN^LPC-KO^ (WT; n = 7) mice. Data are presented as mean ± SEM. ***, p < 0.001; ****, p < 0.0001. (D) Representative hematoxylin/eosin-stained liver section from 10 week-old wild-type (WT) and OTULIN^LPC-KO^ mice demonstrating loss of normal hepatic architecture and multifocal, lobular infiltration of mononuclear cells in the liver of OTULIN^LPC-KO^ mice. Scale bar, 50 μm. (E) Relative mRNA levels of *Il6, Il1b, Mcp1* and *Rantes* in total liver lysates from 10 week-old OTULIN^LPC-KO^ mice (n = 8) and control (WT, n = 11) mice. Data are presented as mean ± SEM. **, p < 0.01; ***, p < 0.001; ****, p < 0.0001. (F-G) Cytokeratin 19 (CK19) (F) and Sirius Red (G) staining on liver sections from 10 week-old wild-type (WT) and OTULIN^LPC-KO^ mice demonstrating oval cell hyperplasia and fibrosis in OTULIN^LPC-KO^ livers. Scale bar, 200 μm. (H) Representative Oil red O stained liver cryosections from 10 week-old wild-type (WT) and OTULIN^LPC-KO^ mice. Scale bar, 200 μm. (I) Representative hematoxylin/eosin-stained liver section showing early well-differentiated HCC in 6 month-old OTULIN^LPC-KO^ mice. T, tumor. Scale bar, 200 μm.

Dissection of livers from 8-10-week-old OTULIN^LPC-KO^ mice revealed the presence of a severe liver phenotype. All OTULIN^LPC-KO^ livers displayed hepatomegaly and an aberrant liver architecture with presence of numerous small but macroscopically visible nodules, in contrast to littermate control mice that did not show any overt liver pathology (Fig. 1B, Suppl. Fig. 2). Alongside, OTULIN^LPC-KO^ mice displayed elevated levels of the liver-specific enzymes aspartate transaminase (AST), alanine transaminase (ALT), and alkaline phosphatase (ALP), indicative of liver damage (Fig. 1C), and also displayed hyperbilirubinemia, indicative of cholestasis (Fig. 1C). Histological analysis confirmed a severe chronic inflammatory liver disease characterized by immune cell infiltration, and associated with hyperplasia and hypertrophy of hepatocytes. In addition, the OTULIN-deficient livers showed multifocal cell death of individual hepatocytes and an increase in the number of mitotic figures and nuclear sizes (polyploidy) (Fig. 1D, Suppl. Fig. 3). FACS analysis further suggested an inflammatory phenotype with an increased proportion of monocytes in OTULIN-deficient livers (Suppl. Fig. 4). In addition, this analysis also identified a significant proportion of Clec4F+Tim4-Kupffer cells (KCs) in OTULIN^LPC-KO^ livers. Typically the presence of Tim4-KCs suggests KC death and their subsequent replacement from circulating monocytes which only gain Tim4-expression with time (Scott et al., 2016) (Suppl. Fig. 4). Increased expression of pro-inflammatory cytokines and chemokines (*Il6, IL1β Rantes* and *Mcp1*) could also be demonstrated in the OTULIN-deficient liver lysates (Fig. 1E). Sirius red staining, *Tgfb1* mRNA expression and CK19 immunostaining confirmed the presence of fibrosis, associated with oval cell hyperplasia, in the livers of OTULIN^LPC-KO^ mice (Fig. 1F-G, Suppl. Fig.5). Finally, OTULIN-deficient livers showed signs of steatosis with increased lipid deposition in hepatocytes (Fig. 1H). In contrast to OTULIN^LPC-KO^ mice, liver sections from wild-type mice showed a normal hepatic parenchyma and no signs of inflammation, steatosis or fibrosis (Fig. 1D-H, Suppl. Fig. 3-5).

The liver phenotype of OTULIN^LPC-KO^ mice, characterized by chronic inflammation, steatosis and fibrosis, resembles non-alcoholic steatohepatitis (NASH) in humans and represents a risk factor for the development of HCC (Sherman, 2005). To evaluate if OTULIN^LPC-KO^ lesions represent pre-neoplastic lesions which may develop at later stages into HCC, livers from 6-month-old OTULIN^LPC-KO^ mice were dissected. All OTULIN^LPC-KO^ mice displayed a severe liver phenotype with presence of multiple nodules (Suppl. Fig. 6A), which were not detected in the liver of control littermate mice. Histology demonstrated the presence of early well-differentiated HCC in three out of eight knockout cases (37,5%) (Fig. 1I), while the other five knockout livers displayed diffuse and severe pre-neoplastic lesions reminiscent of HCC. Malignancy was further confirmed in young OTULIN^LPC-KO^ mice by increased RNA levels of several oncofetal HCC marker genes, such as alpha-fetoprotein (AFP) and connective tissue growth factor (CTGF), as well as of hepatic stem cell marker genes, like glypican 3 (GPC-3) and CD133 (Suppl. Fig. 6B-C). By the age of 1 year, all OTULIN^LPC-KO^ mice had developed severe liver pathology displaying multiple neoplastic lesions, ranging from adenoma to HCC (Suppl. Fig. 6D-G).

### OTULIN deletion leads to accumulation of linear ubiquitin chains but paradoxically causes spontaneous hepatocyte apoptosis

LUBAC-mediated linear ubiquitination plays an important role in the activation of the NF-κB signaling pathway and in protecting cells from death (Peltzer et al., 2014, 2018). OTULIN is reported to counteract LUBAC activity by removing the linear chains conjugated to LUBAC substrates, which include TNFR1, NEMO, RIPK1, RIPK2 and MyD88 (Hrdinka and Gyrd-Hansen, 2017). In accordance, OTULIN deficiency was shown to cause hyper-ubiquitination of signaling proteins, leading to NF-κB hyper-activation and inflammatory signaling (Hrdinka and Gyrd-Hansen, 2017). A recent report, however, showed that OTULIN promotes rather than counteracts LUBAC activity by preventing its auto-ubiquitination, and that knockin mice expressing DUB inactive OTULIN resemble LUBAC-deficient mice, and die at midgestation due to pro-inflammatory cell death (Heger et al., 2018). We could demonstrate that specific deletion of OTULIN in hepatocytes resulted in reduced expression of HOIP and SHARPIN (Fig. 2A), as previously reported in other cell types (Heger et al., 2018; Damgaard et al., 2016, 2019), including MEFs (Suppl. Fig. 1A). Despite the reduced LUBAC expression levels, analysis of linear ubiquitination by specific UBAN pulldowns revealed massive accumulation of linear polyubiquitin in liver lysates of OTULIN-deficient mice (Fig. 2B). Of note, the LUBAC substrate RIPK1 did not show increased linear ubiquitination, suggesting accumulation of linear polyubiquitin on specific substrates or as free chains (Fig. 2B). Surprisingly, the accumulation of linear chains caused by OTULIN deletion correlated with increased apoptosis of hepatocytes. Indeed, in contrast to control livers, OTULIN^LPC-KO^ livers showed dispersed and numerous cleaved caspase-3 and TUNEL positive cells, indicative of apoptosis induction (Fig. 2C-D, Suppl. Fig. 3 and 7). The detection of cleaved caspase-8 and −3 in OTULIN-deficient liver lysates further demonstrated that OTULIN deficiency sensitizes hepatocytes to caspase-dependent extrinsic apoptosis (Fig. 2E).

**Figure 2.**
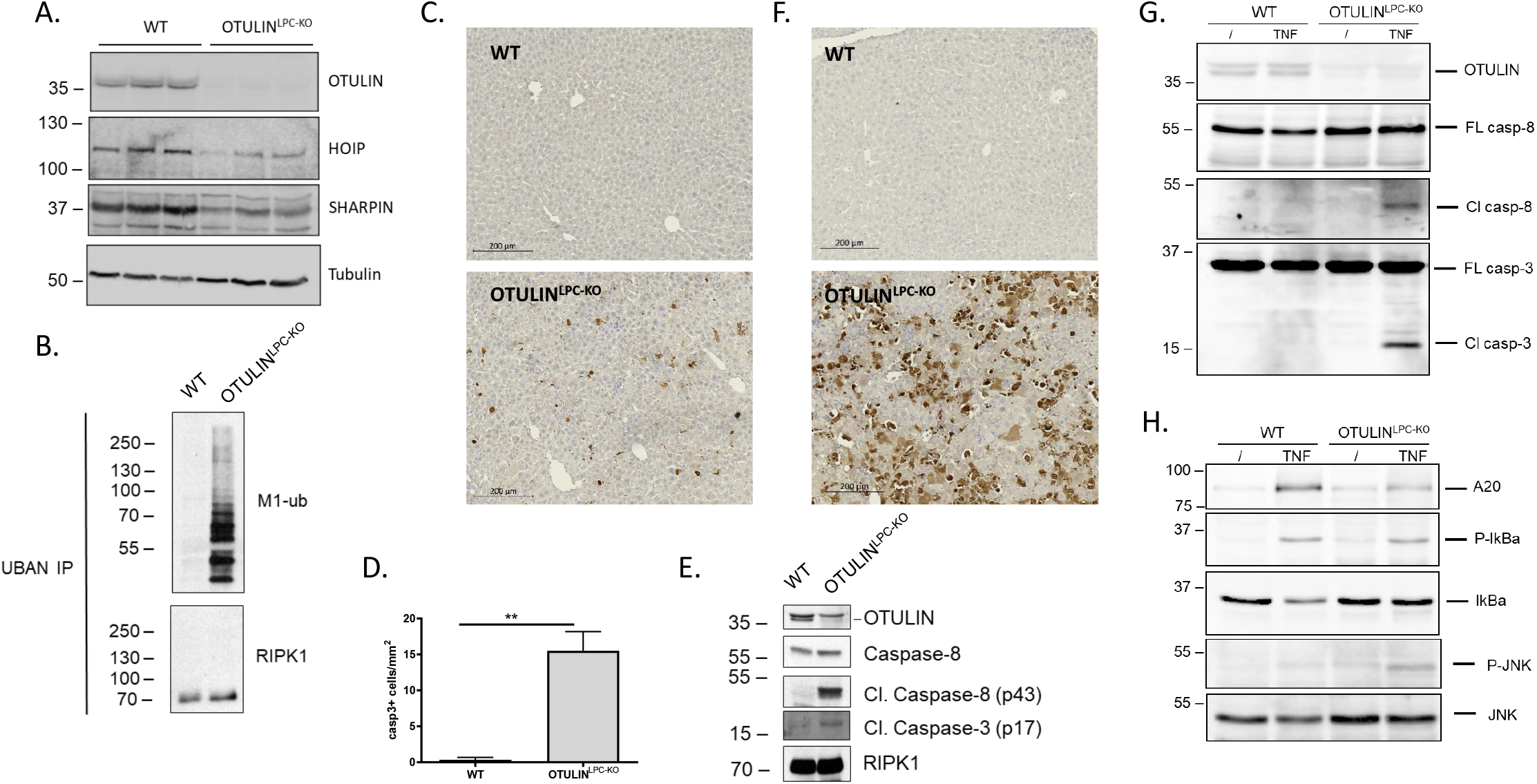
OTULIN suppresses linear ubiquitination and cell death in liver parenchymal cells. (A) Western blot analysis for expression of OTULIN and LUBAC proteins in liver lysates from wild-type (WT) and OTULIN^LPC-KO^ mice. Anti-tubulin immunoblotting was used as loading control. Data are representative of two independent experiments. (B) M1 chains were immunoprecipitated from whole liver cell lysates from untreated OTULIN^LPC-KO^ and control littermate mice (WT) using GST-UBANs. Protein levels were determined by immunoblotting. (C) Representative images of liver sections from 10-week-old OTULIN^LPC-KO^ mice and control (WT) mice after immunostaining for cleaved caspase-3. Scale bar, 200 μm. (D) Quantification of the number of caspase-3 positive cells. Data are presented as mean ± SEM, n = 5 mice per genotype. ** p < 0.01. (E) Western blot analysis for expression of OTULIN, RIPK1 and full-length and cleaved (Cl) caspase-8 and −3 in liver lysates from OTULIN^LPC-KO^ and control littermate mice (WT). Data are representative of three independent experiments. (F) Caspase-3 staining on liver sections from OTULIN^LPC-KO^ mice and control wild-type (WT) littermate mice injected with TNF for 3 h. (G) Western blot analysis for OTULIN, and full-length (FL) and cleaved (Cl) caspase-8 and −3 expression in liver lysates from OTULIN^LPC-KO^ and control littermate mice (WT) either or not injected with 5 μg mouse TNF for 5 h. (H) Western blot analysis for A20, IκBα, phosphorylated IκBα, JNK and phosphorylated JNK in liver lysates from OTULIN^LPC-KO^ and control littermate mice (WT) either or not injected with 5 μg mouse TNF for 5 h.

To better understand the role of OTULIN in protecting hepatocytes from apoptosis, we evaluated the consequences of its deficiency in the TNFR1 signaling pathway. For this, OTULIN^LPC-KO^ mice and control littermate mice were injected with a normally sublethal dose of recombinant mouse TNF (5 μg / 20 g of body weight). In contrast to control livers, OTULIN^LPC-KO^ livers displayed numerous cleaved caspase-3 positive hepatocytes in response to the TNF challenge (Fig. 2F). Hepatocyte apoptosis was further confirmed by the presence of cleaved caspase-8 and −3 in the liver lysates (Fig. 2G). The sensitization to apoptosis was associated with a defect in TNF-induced NF-κB activation, as monitored by reduced IκBα degradation and reduced expression of the NF-κB response protein A20 (Fig. 2H). In contrast, the ratio between total and activated JNK was similar in the liver lysates of OTULIN^LPC-KO^ and control mice upon TNF challenge (Fig. 2H).

Together, these data demonstrate the importance of OTULIN in restricting M1 ubiquitination in hepatocytes, and identify OTULIN as an essential protein protecting mice from hepatocyte apoptosis and acute liver failure. These results also demonstrate that despite spontaneous accumulation of linear polyubiquitin in OTULIN-deficient livers, signaling to NF-κB by TNF is compromised, which may provide a molecular explanation for the spontaneous apoptosis of OTULIN-deficient hepatocytes.

### Hepatocyte apoptosis is accompanied with compensatory hepatocyte proliferation in OTULIN-deficient livers

The liver has remarkable regenerative capacity, and hepatocytes massively start proliferating following hepatic loss in order to restore liver function and mass (Luedde et al., 2014). Since hepatocyte apoptosis in OTULIN deficient mice may be the driving force triggering hepatocyte proliferation favoring hepatocarcinogenesis, we next assessed liver proliferation in OTULIN^LPC-KO^ mice. Indeed, hepatocyte apoptosis in OTULIN^LPC-KO^ mice was accompanied by excessive liver cell proliferation, as demonstrated by increased number of Ki67-positive cells (Fig. 3A-B). In agreement, expression of the cell-cycle markers cyclin D1 and PCNA was strongly enhanced in liver lysates of OTULIN^LPC-KO^ mice (Fig. 3C-D). Together, these data underline the correlation between hepatocyte cell death and proliferation, and suggest that the severe liver pathology in OTULIN^LPC-KO^ mice develops as a result of continuous hepatocyte apoptosis and compensatory hepatocyte proliferation, promoting the development of chronic hepatitis, steatosis, fibrosis and eventually HCC.

**Figure 3.**
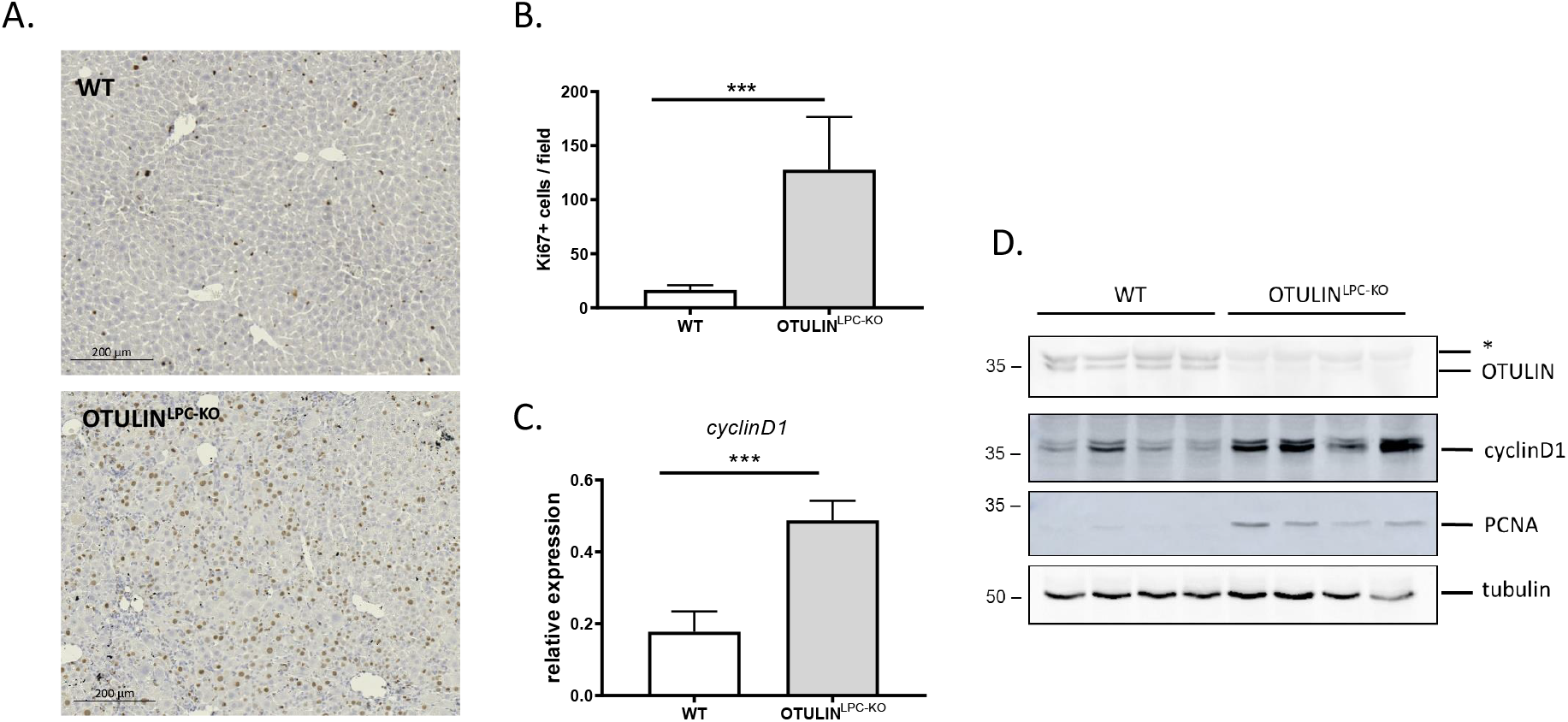
Enhanced hepatocyte proliferation in OTULIN-deficient liver. (A) Representative images of liver sections from 10 week-old OTULIN^LPC-KO^ mice and control (WT) mice after Ki67 immunostaining for proliferating cells. Scale bar, 200 μm. (B) Quantification of the number of Ki67 positive cells. Data are presented as mean ± SEM, n = 5 mice per genotype. *** p < 0.001. (C) Relative mRNA expression of *Cyclin-D1* in total liver lysates from 10 week-old OTULIN^LPC-KO^ mice (n = 12) and control (WT, n = 11) mice. Data are presented as mean ± SEM. ***, p < 0.001. (D) Liver protein extracts from 10 week-old OTULIN^LPC-KO^ mice and control (WT) mice were subjected to Western blotting using antibodies detecting OTULIN, cyclinDl and PCNA. Anti-tubulin immunoblotting was used as loading control. Data are representative of three independent experiments.

### FADD and RIPK1 kinase activity-dependent apoptosis drives hepatocyte death and liver pathology in OTULIN^LPC-KO^ mice

Since OTULIN was previously shown to be essential to prevent TNF-dependent systemic inflammation in humans and in mice (Zhou et al., 2016; Damgaard et al., 2016, 2019), we next addressed the specific role of TNF in triggering the spontaneous death of OTULIN-deficient hepatocytes leading to the development of spontaneous liver pathology in OTULIN^LPC-KO^ mice. For this, we generated TNF deficient OTULIN^LPC-KO^ mice. Surprisingly, genetic deletion of *Tnf* did not prevent spontaneous liver pathology in OTULIN^LPC-KO^ mice (Suppl. Fig. 8A). Although 10-week-old OTULIN^LPC-KO^/TNF^KO^ mice had lower levels of serum ALT, AST and ALP compared to the levels seen in OTULIN^LPC-KO^ mice (Suppl. Fig. 8B), histological analysis of liver tissue revealed that the additional deletion of TNF did not significantly reduce immune cell infiltration or the numbers of caspase-3-positive hepatocytes (Suppl. Fig.8C-E). Similarly, knockout of TNFR1 in all cells did not protect OTULIN^LPC-KO^ mice from developing liver disease (Suppl. Fig. 8A-E). Together, these results demonstrate that TNF signaling does not drive chronic liver damage in OTULIN^LPC-KO^ mice.

Previous studies have demonstrated that the protective role of linear ubiquitination against RIPK1 kinase-dependent and -independent FADD-mediated apoptosis is not limited to TNFR1 signaling (Lafont et al., 2017; Taraborrelli et al., 2018), suggesting conserved regulatory mechanisms downstream of various death receptors. Since RIPK1 has already been implicated in liver pathology by regulating hepatocyte death (Kondylis and Pasparakis, 2019), we first evaluated the presence and consequence of RIPK1 kinase-dependent apoptosis of hepatocytes in our OTULIN^LPC-KO^ mice. To do so, we crossed OTULIN^LPC-KO^ mice to knock-in mice expressing a kinase-inactive RIPK1-D138N mutant (Polykratis et al., 2014). 10-week-old OTULIN^LPC-KO^/RIPK1^D138N/D138N^ mice demonstrated slightly reduced liver pathology but showed significantly reduced serum ALT, AST and ALP levels, and reduced *Tgfb1* expression compared to OTULIN^LPC-KO^ mice (Fig. 4A, B). Although the liver from OTULIN^LPC-KO^/RIPK1^D138N/D138N^ mice still revealed an aberrant tissue architecture, significantly reduced numbers of apoptotic hepatocytes but not of proliferating hepatocytes could be observed in livers from OTULIN^LPC-KO^/RIPK1^D138N/D138N^ mice compared to OTULIN^LPC-KO^ mice (Fig. 4C-E).

**Figure 4.**
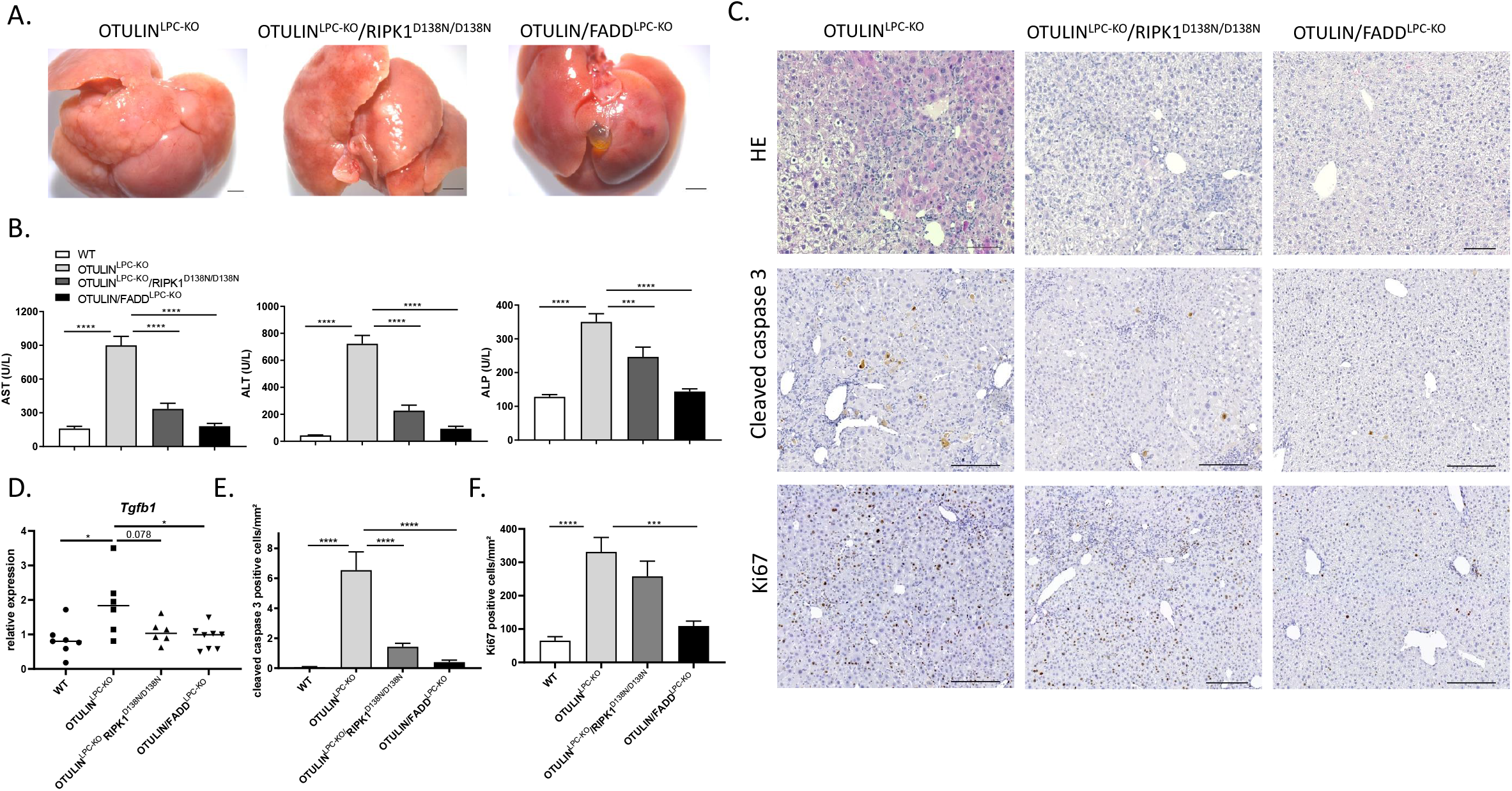
FADD deficiency or absence of RIPK1 kinase activity protects OTULIN^LPC-KO^ mice from developing liver pathology. (A) Macroscopic pictures of representative livers from a 10 week-old OTULIN^LPC-KO^, OTULIN^LPC-KO^/RIPK1^D138N/D138N^ and OTULIN/FADD^LPC-KO^ mouse. Scale bar, 2 mm. (B) Serum ALT, AST and ALP levels in control (WT, n = 49), OTULIN^LPC-KO^ (n = 22), OTULIN^LPC-KO^/RIPK1^D138N/D138N^ (n = 11) and OTULIN/FADD^LPC-KO^ (n = 18) mice. Data are presented as mean ± SEM. ***, p < 0.001; ****, p < 0.0001. (C) Representative hematoxylin/eosin, cleaved caspase 3 and Ki67-stained liver section from 10 week-old OTULIN^LPC-KO^, OTULIN^LPC-KO^/RIPK1^D138N/D138N^ and OTULIN/FADD^LPC-KO^ mice. Scale bar H&E, 100 μm, cleaved caspase 3 and Ki67, 200 μm. (D) Relative mRNA expression of *Tgfb1* in total liver lysates from 10 week-old control (WT, n = 7), OTULIN^LPC-KO^ mice (n = 6), OTULIN^LPC-KO^/RIPK1^D138N/D138N^ (n=6) and OTULIN/FADD^LPC-KO^ mice (n=8) mice. Data are presented as mean ± SEM. * < 0,05. (E) Quantification of the number of cleaved caspase-3 positive cells in liver sections from control (WT, n = 6), OTULIN^LPC-KO^ (n = 5), OTULIN^LPC-KO^/RIPK1^D138N/D138N^ (n = 7) and OTULIN/FADD^LPC-KO^ (n = 8) mice. Data are presented as mean ± SEM. ****, p < 0.0001. (F) Quantification of the number of Ki67 positive cells in liver sections from control (WT, n = 9), OTULIN^LPC-KO^ (n = 6), OTULIN^LPC-KO^/RIPK1^D138N/D138N^ (n = 8) and OTULIN/FADD^LPC-KO^ (n = 7) mice. Data are presented as mean ± SEM. ***, p < 0.001; ****, p < 0.0001.

Because RIPK1 kinase activity can induce both FADD-dependent apoptosis and RIPK3/MLKL-dependent necroptosis (Pasparakis and Vandenabeele, 2015), we next evaluated occurrence of MLKL-dependent hepatocyte necroptosis. OTULIN^LPC-KO^ mice were crossed to mice with a floxed *Mlkl* allele (Murphy et al., 2013), generating mice lacking both OTULIN and MLKL specifically in liver parenchymal cells. MLKL deficiency could, however, not prevent liver pathology in OTULIN^LPC-KO^ mice, as shown by liver histology and detection of liver damage in OTULIN/MLKL^LPC-KO^ mice (Suppl. Fig. 8), arguing against a role for MLKL-driven necroptosis in the OTULIN^LPC-KO^ pathology.

To confirm that hepatocyte apoptosis is responsible for the development of liver pathology in OTULIN^LPC-KO^ mice, we next generated OTULIN^LPC-KO^ mice lacking FADD specifically in liver parenchymal cells by crossing the OTULIN^LPC-KO^ line with mice having a floxed *Fadd* allele (FADD^FL/FL^) (Guire et al., 2010), thereby preventing both RIPK1 kinase-dependent and - independent FADD-mediated apoptosis. In contrast to OTULIN^LPC-KO^ mice, which all developed severe liver pathology, OTULIN/FADD^LPC-KO^ mice had a completely normal tissue architecture and did not display liver damage, as detected by tissue histology, by serum analysis of ALT, AST and ALP levels, and by expression of *Tgfb1* reminiscent of liver fibrosis (Fig. 4A-D). In agreement, no hepatocyte cell death, shown by cleaved caspase-3 detection in liver tissue sections, could be detected in livers from OTULIN/FADD^LPC-KO^ mice (Fig. C, E). In addition to rescuing hepatocyte death, FADD deficiency also inhibited the increased proliferation observed in the liver of OTULIN^LPC-KO^ mice, as shown by the reduced Ki67 staining in liver lysates from OTULIN/FADD^LPC-KO^ mice (Fig. 4C, F). This demonstrates that the increased hepatocyte proliferation and inflammation in liver of OTULIN^LPC-KO^ mice is a secondary response to the apoptosis of OTULIN deficient hepatocytes. Collectively, these data show that liver disease in OTULIN^LPC-KO^ mice develops as a consequence of FADD-dependent apoptosis, partially driven by RIPK1 kinase activity, but not necroptosis, of OTULIN deficient hepatocytes.

Finally, since inflammation-induced liver injury is often associated with increased intestinal permeability and bacterial translocation to the liver via the portal vein (Seki and Schnabl, 2012), we assessed the importance of MyD88-dependent signaling for the development of liver pathology in OTULIN^LPC-KO^ mice. 10-week-old OTULIN^LPC-KO^/MyD88^KO^ mice were, however, not protected from developing liver disease and all exhibited similar increased serum ALT, AST and ALP levels compared with OTULIN^LPC-KO^ mice (Suppl. Fig. 8A-B). Also on histology, no differences in immune cell infiltration, hepatocyte cell death and proliferation could be observed (Suppl. Fig. 8C-E).

### Interferon signaling contributes to the liver pathology in hepatocyte-specific OTULIN knockout mice

The liver phenotype of OTULIN^LPC-KO^ mice was associated with a significant upregulation of inflammatory gene expression, including the expression of interferon response genes, which were all reverted to baseline levels in the FADD deficient genetic background (Fig. 1E, Fig. 5A-B). This suggests that OTULIN might also be important to suppress the production of type I interferons, as suggested by recent studies demonstrating the promotion of IFNAR signaling in conditions where OTULIN or LUBAC signaling is impaired (Heger et al., 2018; Peltzer et al., 2018). To investigate if type I interferon signaling contributes to the development of liver pathology in OTULIN^LPC-KO^ mice, we generated OTULIN^LPC-KO^ mice lacking the interferon-α receptor 1 (IFNAR1) (Muller et al., 1994). 10-week-old OTULIN^LPC-KO^/IFNAR1^KO^ mice demonstrated reduced liver pathology and showed significantly reduced AST, ALT and ALP levels compared to OTULIN^LPC-KO^ mice (Fig. 5C-D). In addition, significantly reduced numbers of apoptotic hepatocytes but not of proliferating hepatocytes could be observed in livers from OTULIN^LPC-KO^/IFNAR1^KO^ mice compared to OTULIN^LPC-KO^ mice (Fig. 5E-G). Together these data demonstrate that type I interferons are produced in OTULIN deficient liver and critically contribute to the severe liver phenotype in OTULIN^LPC-KO^ mice.

**Figure 5.**
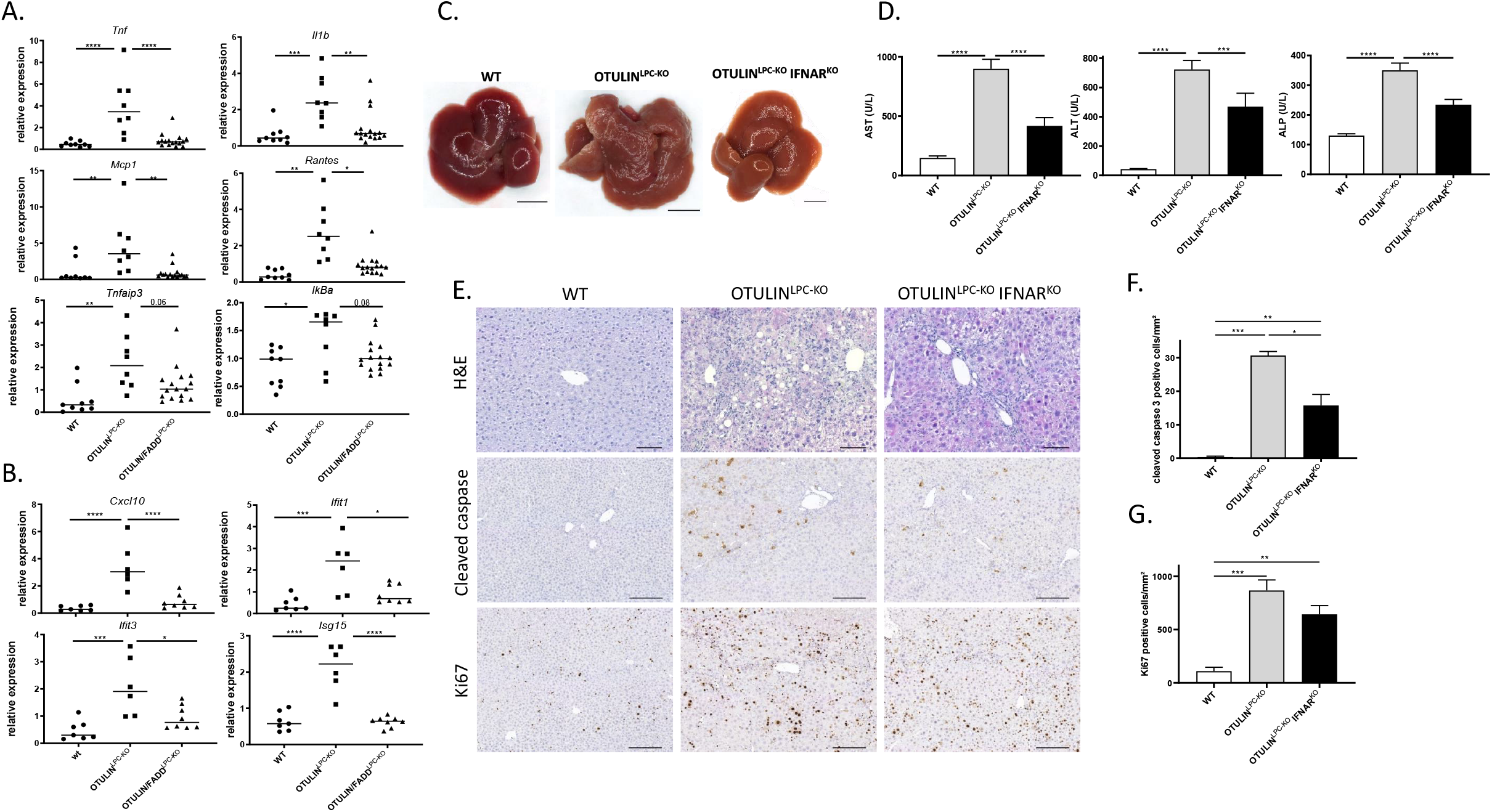
Interferon signaling contributes to the liver pathology in OTULIN^LPC-KO^ mice. (A) Relative mRNA expression of *Tnf, Il1b, Mcp1, Rantes, Tnfaip3* and *IkBa* in total liver lysates from 10 week-old control (WT, n = 9), OTULIN^LPC-KO^ mice (n = 8) and OTULIN/FADD^LPC-KO^ mice (n = 16) mice. (B) Relative mRNA expression of *Cxcl10, Ifit1, Ifit3* and *Isg15* in total liver lysates from 10 week-old control (WT, n = 7), OTULIN^LPC-KO^ mice (n = 6) and OTULIN/FADD^LPC-KO^ mice (n = 8) mice. Data are presented as mean ± SEM. *, p< 0,05; **, p < 0.01; ***, p < 0.001; ****, p < 0.0001. (C) Macroscopic pictures of representative livers from a 10 week-old control (WT), OTULIN^LPC-KO^ and OTULIN^LPC-KO^/IFNAR1^KO^. Scale bar, 5 mm. (D) Serum ALT, AST and ALP levels in control (WT, n = 57), OTULIN^LPC-KO^ (n = 22), and OTULIN^LPC-KO^/IFNAR1^KO^ (n=12) mice. Data are presented as mean ± SEM. ***, p < 0.001; ****, p < 0.0001. (E) Representative hematoxylin/eosin, cleaved caspase 3 and Ki67-stained liver section from 10 week-old control (WT), OTULIN^LPC-KO^ and OTULIN^LPC-KO/^IFNAR1^KO^. Scale bar H&E, 100 μm, cleaved caspase 3 and Ki67, 200 μm. (F) Quantification of the number of cleaved caspase-3 positive cells in liver sections from control (WT, n = 4), OTULIN^LPC-KO^ (n = 3) and OTULIN^LPC-KO^/IFNAR^KO^ mice (n = 6). Data are presented as mean ± SEM. *, p < 0.05; ** p < 0.01; ***, p < 0.001. (G) Quantification of the number of Ki67 positive cells in liver sections from control (WT, n = 4), OTULIN^LPC-KO^ (n = 3) and OTULIN^LPC-KO^/IFNAR^KO^ mice (n = 6). Data are presented as mean ± SEM. ** p < 0.01; ***, p < 0.001

## Discussion

The incidence of hepatitis and HCC has risen in Western countries, most probably because of changes in dietary habits causing metabolic stress, the so called ‘metabolic syndrome’, and non-alcoholic fatty liver disease (NAFLD). HCC develops as a result of a chronic liver inflammation, and is mostly diagnosed at advanced stages with very limited treatment options. Hence, early and sustained suppression of chronic liver damage is key to reduce the risk to develop HCC (Ringelhan et al., 2018). We here demonstrated the spontaneous TNF-independent but FADD-dependent apoptosis of hepatocytes as the crucial event driving liver inflammation, fibrosis and HCC in hepatocyte-specific OTULIN deficient mice. Our study also demonstrated that RIPK3-MLKL-dependent hepatocyte necroptosis is not involved in the pathology of OTULIN^LPC-KO^ mice, likely due to the absence of RIPK3 expression in hepatocytes, as previously demonstrated (Krishna-Subramanian et al., 2019). Intriguingly, OTULIN/MLKL^LPC-KO^ mice, lacking both OTULIN and MLKL only in the hepatocytes, seem to display an even worse liver phenotype compared to OTULIN^LPC-KO^ mice. This would suggest that MLKL-dependent necroptosis, eventually through a pathway independent of RIPK3, would protect OTULIN deficient hepatocytes from death by apoptosis. Interestingly, a recent study demonstrated that suppression of necroptosis in hepatocytes could promote hepatocyte apoptosis favoring the development of HCC (Seehawer et al., 2018). This observation suggested that hepatocyte necroptosis generates a liver cytokine microenvironment which promotes the development of intrahepatic cholangiocarcinoma and not HCC from oncogenically transformed hepatic cells (Seehawer et al., 2018). More studies will be needed to investigate this further.

The protective role of LUBAC-mediated linear ubiquitination is well-established downstream of TNFR1. On the one hand, it promotes the transcriptional upregulation of anti-apoptotic proteins by NF-κB, which protect cells from RIPK1 kinase-independent apoptosis (Wang et al., 2008). On the other hand, it allows the phosphorylation of RIPK1 by IKKα/β- and TBK1/IKKε, which was shown to protect cells from RIPK1 kinase-dependent apoptosis (Dondelinger et al., 2015, 2019; Lafont et al., 2018; Xu et al., 2018; Ting and Bertrand, 2016). While several studies have reported the importance of proper TNF signaling for the maintenance of liver homeostasis and to prevent liver inflammation and inflammation-induced HCC (Luedde et al., 2014), our study did not identify TNF as a driving cytokine in the liver pathology of OTULIN^LPC-KO^ mice. The molecular mechanism by which linear ubiquitination protects cells from death may however not be limited to TNFR1 but instead be conserved between death receptors. Indeed, LUBAC was also shown to protect cells from RIPK1 kinase-dependent and -independent FADD-mediated apoptosis downstream of other TNFR superfamily members (Lafont et al., 2017; Taraborrelli et al., 2018). In accordance with this notion, we found that transgenic expression of a kinase-inactive RIPK1^D138N^ mutant could significantly ameliorate, however not fully rescue, the liver disease. This indicates that hepatocyte apoptosis in OTULIN^LPC-KO^ mice is FADD/caspase-8 dependent, and partially driven by RIPK1 kinase activity. Despite massive accumulation of linear polyubiquitin, signaling to NF-κB by TNF was compromised in OTULIN deficient hepatocytes, suggesting that OTULIN deficiency also affects proper signaling to NF-κB downstream of the receptor that triggers apoptosis in OTULIN^LPC-KO^ mice. Therefore, a defect in the NF-κB-mediated upregulation of cytoprotective mechanisms in OTULIN deficient hepatocytes may account for the RIPK1 kinase-independent hepatocyte apoptosis, while inappropriate regulation of RIPK1 by IKKs may account for the RIPK1 kinase-dependent hepatocyte apoptosis, together inducing liver inflammation and a partial liver pathology.

Since IFNAR1 deficiency protects OTULIN^LPC-KO^ mice, at least partially, from developing severe liver disease, we further demonstrate an important contribution of type I interferons to the phenotype of OTULIN^LPC-KO^ mice. This observation suggests that OTULIN might also suppress pathways responsible for the production of interferons, either indirectly by preventing overall inflammation, or by direct control of interferon production. In that context, FADD and RIPK1 have been identified as being implicated in an immune defense pathway against intracellular dsRNA (Balachandran et al., 2004; Michallet et al., 2008; Rajput et al., 2011; Ingram et al., 2019). Further studies are required to identify the pathway(s) regulated by OTULIN driving type I interferon production.

Our results using hepatocyte-specific OTULIN deficient mice also confirm the importance of linear ubiquitination and its regulation by OTULIN for the protection against hepatocyte apoptosis and liver inflammation, in agreement with a previous study demonstrating the importance of LUBAC-mediated linear ubiquitination for liver physiology preventing hepatocyte apoptosis, hepatitis and HCC development (Shimizu et al., 2017).

With our studies, we identified OTULIN as a critical regulator of liver homeostasis by protecting the liver parenchymal cells from death by apoptosis responsible for liver inflammation, fibrosis and eventually liver cancer, similar to what has been demonstrated for NEMO (Luedde et al., 2007; Kondylis et al., 2015). Blockade of hepatocyte cell death may thus have therapeutic potential for patients suffering from chronic inflammatory liver diseases. Also, hepatocyte-specific OTULIN deficient mice may serve as a novel model to study NAFLD and HCC and help to develop new treatments for the diagnosis or therapeutic intervention in patients suffering from inflammation-induced liver pathologies risking the development of HCC.

## Materials and Methods

### Mice

The following mouse lines were used: Alfp-Cre (Kellendonk et al., 2000), *Fadd*^FL^ (Guire et al., 2010), *Mlkl*^FL^ (Murphy et al., 2013), *Ripk1*^D138N^ (Polykratis et al., 2014), *Myd88*^-/-^ (Adachi et al., 1998), *Tnf*^-/-^ (Pasparakis et al., 1996), *Tnfr1*^−/−^ (Pfeffer et al., 1993) and *Ifnar1*^−/−^ (Muller et al., 1994). All alleles were maintained on a C57BL/6 genetic background. Littermates carrying the floxed alleles but not the Alfp-Cre transgene served as controls. All experiments were performed on mice with C57BL/6 genetic background. Mice were housed in individually-ventilated cages at the VIB Center for Inflammation Research, in a specific pathogen-free animal facility. All experiments on mice were conducted according to institutional, national and European animal regulations. Animal protocols were approved by the ethics committee of Ghent University.

### Generation of hepatocyte-specific OTULIN knockout mice

C57BL/6 embryonic stem (ES) cells with an *Otulin*/*Fam105b^tm1a^* allele were generated by the European Mouse Mutagenesis (EUCOMM) Programme using a ‘knockout-first with conditional potential’ gene targeting cassette, and used to generate mice bearing a neomycin-LacZ cassette and a loxP-flanked exon 3 of *Otulin*. For this, the targeted ES cell clone was injected into 3.5-day blastocysts and transferred to the uteri of pseudopregnant foster mothers. Male chimeras were mated with C57BL/6 females to obtain germline transmission of the *Otulin* floxed allele (still containing the neomycine selection cassette, *Otulin*^NFL^). The Frt-flanked neomycin-LacZ cassette was removed by crossing *Otulin^NFL^* mice with a Flp-deleter strain (Rodríguez et al., 2000) generating an *Otulin* floxed allele (*Otulin*^FL^). *Otulin*^FL/FL^ mice were crossed to Alfp-Cre transgenic mice(Kellendonk et al., 2000) to generate liver parenchymal cell-specific OTULIN knockout (*Otulin*^LPC-KO^) mice (Suppl. Fig. 1).

### Isolation and immortalization of mouse embryonic fibroblasts (MEFs)

Otulin^+/−^ mice were generated by crossing chimeras transmitting the *Otulin*^FL^ genotype with Cre-deleter mice. MEFs were prepared from E10.5 Otulin^+/+^, Otulin^+/−^ and Otulin^-/−^ embryos, and immortalized through serial passaging and frozen in liquid nitrogen

### Isolation of liver cells for immune profiling

The protocol for the isolation of liver cells was adapted from Mederacke et al (Mederacke et al., 2015) as previously described (Bonnardel et al. Immunity, in press). Briefly, after retrograde cannulation, livers were first perfused with an EGTA-containing solution for 1-2 minutes following by a perfusion with 0.2 mg/ml collagenase A-containing solution for 5 minutes (6 ml/min). Livers were minced and incubated for 20 minutes with 0.4 mg/ml Collagenase A and 10 U/ml DNase in a water bath at 37°C. All subsequent procedures were performed at 4°C. After filtration with a 100 μm mesh filter, the cell suspensions were centrifuged at 400 g for 7 min and re-suspended in 2 ml of red blood lysis buffer for 3 minutes. Suspensions were washed in PBS and further filtered on a 40 μm mesh filter and centrifuged twice for 1 min at 50 g resulting in an hepatocytes-enriched fraction (pellet) and a leukocytes/LSECs/HSCs-enriched fraction (supernatant). Both fractions were further centrifuged at 400 g for 7 min before proceeding with FACS staining. After staining cells were analysed on a BDSymphony and resulting data were analysed using FlowJo Software. Following antibodies were used: CD26-FITC (BD, Cat no: 559652), CD172a-BB630P (BD, Custom conjugate), Tim4-Percp-eFluor710 (Thermo Fischer, Cat no: 46-5866-82), Clec4F-unconjugated (R&D systems, Cat no: AF2784), Goat IgG-AF647 (Thermo Fischer, Cat no: A-21447), Fixable Live/dead dye – APCeFluor780, Ly6C-eFluor450 (eBioscience – Cat no: 48-5932-82), CD45-BV510 (Biolegend, Cat no: 103138), CDllb-BV605 (BD, Cat no: 563015), CD64-BV711 (Biolegend, Cat no: 139311), F4/80-BV786 (Biolegend, Cat no: 123141), XCR1-BV650 (Biolegend, Cat no: 148220), SiglecF-BUV395 (BD, Cat no: 740280), Ly6G-BUV563 (BD, Cat no: 612921), MHCII-BUV805 (BD, custom conjugate), CD11c-PE-eFluor610 (eBioscience, Cat no: 61-0114-82), Lineage markers – CD3 (TONB55-0031-U100, Tonbo Biosciences), CD19 (15-1093-83, eBioscience), B220 (553091, BD), NK1.1-PE-Cy5 (108716, Biolegend).

### Western blot analysis

Cells and liver tissue were homogenized using E1A lysis buffer (50 mM HEPES pH7.6; 250 mM NaCl; 5 mM EDTA; 0.5% NP40) and RIPA (50 mM Tris-HCl, pH 7.6; 1 mM EDTA; 150 mM NaCl; 1% NP-40; 0.5% sodiumdeoxycholate; 0.1% SDS) buffer respectively containing protease inhibitors (Roche) and phosphatase inhibitors (Sigma), denaturated in 1 x Laemmli buffer (50 mM Tris-HCl pH8.2; 2% SDS; 10% glycerol; 0.005% BFB; 5% β-mercapto-ethanol) and boiled for 10 min at 95 °C. 50 μg lysates were separated by SDS-polyacrylamide gel electrophoresis (PAGE), transferred to nitrocellulose and analyzed by immunoblotting. Membranes were probed with antibodies against OTULIN (1:1000, Cell Signaling Technology), A20 (1:1000, Santa Cruz Biotechnologies), caspase-3 (full length and cleaved) (1:1000, Cell Signaling Technology), JNK (1:1000, BD Bioscience), phospho-JNK (1:1000, Sigma-Aldrich), IκBα (1:1000, Santa Cruz Biotechnologies), phospho-IκBα (1:1000, Cell Signaling Technology), HOIL-1 (1:2000, kind gift of Dr. Henning Walczak, UCL London), HOIP (1:1000, kind gift of Dr. Rune Damgaard, MRC Cambridge), Sharpin (1:1000, Proteintech, 1:1000), PCNA (1:1000, Novus), cyclin-D1 (1:1000, Abcam), tubulin (1: 1000, Sigma-Aldrich) and actin-HRP (1:10000, MP Biomedicals). As secondary antibodies, HRP coupled anti-rabbit-HRP, anti-mouse-HRP and anti-goat-HRP were used (1:2500, Amersham) and detection was done by chemiluminescence (Western Lightning Plus ECL, Perkin Elmer) using the Amersham Imager 600 (GE Healthcare).

### Immunoprecipitation

Recombinant GST-UBAN was produced in BL21(DE3) cells. In brief, BL21(DE3) cells were transformed with the plasmid encoding GST-UBAN and protein expression was induced with 0.5 M IPTG. After 4 hours, cells were collected and lysed in lysis buffer (20 mM Tris-HCl pH 7.5, 10 mM EDTA, 5mM EGTA, 150 mM NaCl, 1mM DTT supplemented with phosphatase and protease inhibitor cocktail tablets (Roche Diagnostics)), sonicated and cleared by centrifugation. After centrifugation, Triton-X100 (0.5 % final concentration) was added to the supernatant, which was then transferred onto prewashed glutathione beads and left rotating for 2 hours at 4°C. After incubation, the beads were centrifuged, washed twice with washing buffer (20 mM Tris-HCl pH 7.5, 10 mM EDTA, 150 mM NaCl, 0.5% Triton-X100) and resuspended in resuspension buffer (20 mM Tris-HCl pH 7.5, 0.1% β-mercaptoethanol, 0.05% sodiumazide), ready to be used. Cell lysates from total liver tissue were prepared as described before, protein concentration was determined and 5 mg protein lysate was incubated overnight with GST-UBAN-containing glutathione beads. The next day, the beads were washed three times in RIPA lysis buffer (150 mM NaCl, 1% NP-40, 0.5% Sodium Deoxycholate, 0.1% SDS, 10 mM Tris-HCl pH 8 supplemented with phosphatase and protease inhibitor cocktail tablets (Roche Diagnostics)). Beads were then resuspended in 60μL 1x laemmli buffer for direct analysis.

### Liver injury

Mice were injected i.p. with a sublethal dose of mouse TNF (5 μg mouse TNF/20 g body weight). E. coli-derived recombinant mTNF had a specific activity of 9.46 x 107 IU/mg, and was produced and purified to homogeneity in our laboratory with endotoxin levels not exceeding 1 ng/mg protein. Mice were euthanized after 2.5 hours for histological analysis.

Serum alkaline phosphatase (ALP), alanine transaminase (ALT), aspartate transaminase (AST) and bilirubin levels were measured in the Laboratory of Clinical Biology of the University Hospital Gent, according to standard procedures.

### Histology

Livers were dissected and fixed in 4 % paraformaldehyde and embedded in paraffin. Sections were stained with Hematoxylin/Eosin (H&E) or with specific stains or antibodies. For immunohistochemistry, following dewaxing, slides were incubated in antigen retrieval solution (Dako), boiled for 20 min in a PickCell cooking unit and cooled down for 3 h. Endogenous peroxidase activity was blocked by incubating the slides in 3% wv H2O2 (Sigma). The blocking buffer contains 0.2% goat serum, 0.5% Fish skin gelatin and 2% BSA in PBS. Subsequently slides were incubated with primary antibodies overnight at 4 °C: anti-cleaved caspase 3 (1:300, Cell Signaling) or anti-Ki67 (1:1000, Cell Signaling). Next, the slides were incubated with biotin coupled goat anti-rabbit (1:500, Dako). Subsequently, slides were incubated in ABC solution (Vectastain ELITE ABC Kit Standard, Vector Laboratories). Peroxidase activity was detected by adding diaminobutyric acid (DAB) substrate (ImmPACT DAB Peroxidase (HRP) Substrate, Vector Laboratories), and slides were counterstained with hematoxylin (Sigma), dehydrated and mounted with Entellan (Merck). The pictures were taken with the Zeiss slide scanner. For Oil Red O staining, liver samples were fixed in 4% PFA for 2 hours and incubated in 30 % sucrose overnight after which they were frozen in OCT cryostat embedding medium (Neg −50 Kryo-Medium, Thermo Fisher Scientific). Coupes were fixed in 4% PFA and subsequently incubated in 100% propylene glycol, Oil Red O (at 60 °C) and 85% propylene glycol. The sections were counterstained with hematoxylin (Sigma).

### Histological scoring

Histological diagnosis was performed on H&E-stained slides using standard histopathological analysis based on cellular morphology of mouse liver lesions (Thoolen et al., 2010). The main lesions observed were classified and eventually scored (0: none/minimal; 1: mild; 2:moderate; 3: severe) as: foci of necrosis, portal and lobular inflammation/immune cells infiltration, anisocytosis and anisokaryosis (respectively, variations of cellular and nuclear size and shape), pigmented macrophages, intranuclear pseudoinclusions, foci of vacuolar changes, lobular dysplasia, hepatocellular hypertrophy, foci of cellular alteration (FCA), adenoma, hepatocellular carcinoma (HCC). Oval cell proliferation was evaluated based on a grading system applied to stage precirrhotic/cirrhotic lesions in mouse liver (Kleiner et al., 2005; Hübscher, 2006). The number of mitotic figures and apoptotic cells was evaluated in 5 randomly selected HPF (high power fields, 40X). Differentiation between dysplastic nodules (DNs) and early HCC was based on criteria published for the analysis of human liver (Kondo, 2009; Park, 2011; Schlageter et al., 2014). Due to the difficulty in distinguishing early HCC from DNs in some of the liver samples, a further category indicated with “indiscernible” nodular lesion has been introduced, characterized by a transitional histological appearance in between the two lesions. The immunohistochemistry for Ki67 and CK19 was also evaluated in order to distinguish between these two lesions. The following criteria have been applied to analyze the nodules: 1) higher cellularity (usually 2 times that of the surrounding hepatic parenchyma in HCC); 2) plate thickness and/or pseudoglandular (microacinar) structures (typical of HCC); 3) stromal invasion: well differentiated tumour cells invade the fibrous tissue of portal tracts with lack of ductular reaction (evident by the analysis of CK19 stained slides, typical of HCC); 4) presence of portal tracts (evident by CK19 IHC, present in varying number, usually reduced in HCC; may be present in HNs and DNs); 5) higher number of Ki67-positive hepatocytes (evaluated by IHC, compared to the surrounding parenchyma); 6) vacuolar changes (common in HCC, may be present in DNs); 7) lack of distinct demarcation (more typical of HCC); 8) cellular pleomorphism and loss of normal lobular architecture (present in both DNs and HCC); 9) compression (typical of adenoma when compression of adjacent normal hepatocytes is seen at least on two quadrants). Ki67, Caspase 3 and TUNEL stained slides were evaluated under a microscope, counting the number of positive nuclei in ten randomly selected high-power fields. The total number of positive nuclei for each was divided by 10 to obtain the index. Positive cells within inflammatory foci and areas of oval cell hyperplasia/fibrosis were excluded from the count. The extent of the hepatic parenchyma stained with CK19 and Sirius red stains was evaluated by semiquantitative grading (0: none/minimal; 1: mild; 2: moderate; 3: severe).

### Quantitative real-time PCR

Total RNA was isolated using TRIzol reagent (Invitrogen) and Aurum Total RNA Isolation Mini Kit (Biorad), according to manufacturer’s instructions. Synthesis of cDNA was performed using Sensifast cDNA Synthesis Kit (Bioline) according to the manufacturer’s instructions. cDNA was amplified on quantitative PCR in a total volume of 5 μl with SensiFAST SYBR^®^ No-ROX Kit (Bioline) and specific primers on a LightCycler 480 (Roche). The reactions were performed in triplicates. The following mouse-specific primers were used: *actin* forward GCTTCTAGGCGGACTGTTACTGA; *actin* reverse GCCATGCCAATGTTGTCTCTTAT; *HMBS* forward GAAACTCTGCTTCGCTGCATT; *HMBS* reverse TGCCCATCTTTCATCGTATG; *GAPDH* forward TGAAGCAGGCATCTGAGGG; *GAPDH* reverse CGAAGGTGGAAGAGTGGGAG; *Rpl13a* forward, CCTGCTGCTCTCAAGGTT; *Rpl13a* reverse, TGGTTGTCACTGCCTGGTACTT; *Eef1a1* forward, TCGCCTTGGACGTTCTTTT; *Eef1a1* reverse, GTGGACTTGCCGGAATCTAC; *Cox4i1* forward, AGAATGTTGGCTTCCAGAGC; *Cox4i1* reverse, TTCACAACACTCCCATGTG; *Matrin3* forward, TGGACCAAGAGGAAATCTGG; *Matrin3* reverse, TGAACAACTCGGCTGGTTTC; *TNF* forward, ACCCTGGTATGAGCCCATATAC; *TNF* reverse, ACACCCATTCCCTTCACAGAG; *IL1β* forward, CACCTCACAAGCAGAGCACAAG; *IL1β* reverse, GCATTAGAAACAGTCCAGCCCATAC; *IL6* forward, GAGGATACCACTCCCAACAGACC; *IL6* reverse, AAGTGCATCATCGTTGTTCATACA; *MCP1* forward, GCATCTGCCCTAAGGTCTTCA; *MCP1* reverse, TGCTTGAGGTGGTTGTGGAA; *Rantes* forward, CGTCAAGGAGTATTTCTACAC; *Rantes* reverse, GGTCAGAATCAAGAAACCCT; *Tnfaip3* forward, AAACCAATGGTGATGGAAACTG; *Tnfaip3* reverse, GTTGTCCCATTCGTCATTCC; *IκBα* forward, GTAACCTACCAAGGCTACTC; *IκBα* reverse, GCCACTTTCCACTTATAATGTC; *AFP* forward, CTCAGCGAGGAGAAATGGTC; *AFP* reverse, GAGTTCACAGGGCTTGCTTC; *CTGF* forward, GCCCTAGCTGCCTACCGACT; *CTGF* reverse, GCCCATCCCACAGGTCTTAGA; *GPC3* forward, CTGAGCCGGTGGTTAGCC; *GPC3* reverse TCACTT TCACCATCCCGTCA; *CD133* forward, TGGAGCTACCTGCGGTTTAGA; *CD133* reverse, GGACCTGTGATTGCGATAATGA; *CyclinD1* forward, GCCGAGAAGTTGTGCATCTAC; *CyclinD1* reverse, GGAGAGGAAGTGTTCGATGAA; *Cxcl10* forward, CCAAGTGCTGCCGTCATTTTC; *Cxcl10* reverse, GGCTCGCAGGGATGATTTCAA; *Ifi1* forward, CTACCACCTTTACAGCAACC; *Ifit1* reverse, AGATTCTCACTTCCAAATCAGG; *Ifit3* forward, GCAGCACAGAAACAGATCAC; *Ifit3* reverse, CATTGTTGCCTTCTCCTCAG; *Isg15* forward, ACGGTCTTACCCTTTCCAGTC; *Isg15* reverse, CCCCTTTCGTTCCTCACCAG; *Tgfb1* forward, GCTGAACCAAGGAGACGGAATA; *Tgfb1* reverse, GAGTTTGTTATCTTTGCTGTCACAAGA.

### Statistics

Results are expressed as the mean ± SEM. Statistical significance between WT and OTULIN^LPC-KO^ was assessed using an unpaired two-sample Student t-Test. Statistical significance between OTULIN^LPC-Ko^ and the different genetic crosses was assessed using an one-way ANOVA test, followed by Tukey’s multiple comparison test. The analysis was performed with Prism software.

## ACKNOWLEDGEMENTS

We thank the EUCOMM Consortium for *Otulin*-targeted ES cells. We thank Alexander Warren and James Murphy (The Walter and Eliza Hall Institute of Medical Research, Melbourne, Australia) for the use of floxed *Mlkl* mice. We are grateful to Laetitia Bellen for animal care. L. Verboom is a predoctoral fellow supported by a doctoral scholarship from the Special Research Fund of the Ghent University, and A. Martens is a predoctoral fellow supported by a grant from the “Concerted Research Actions” (GOA) of the Ghent University. Research in the G. van Loo lab is supported by VIB and research grants from the FWO, the “Geneeskundige Stichting Koningin Elisabeth” (GSKE), the CBC Banque Prize, the Charcot Foundation, the “Belgian Foundation against Cancer” and “Kom op tegen Kanker”.

## AUTHOR CONTRIBUTIONS

L.V., A.M., D.P., M.S., H.V., S.V., L.B., C.S. performed the experiments. L.V., E.H., C.S., A.d.B., M.B. and G.v.L. analysed the data. M.P. provided mice. G.v.L. provided ideas and coordinated the project. L.V., M.B. and G.v.L. wrote the manuscript.

## Supplementary Figure legends

**Supplementary Figure 1.** (A) Western blot analysis for expression of OTULIN and LUBAC proteins in wild-type (WT) and OTULIN knockout (KO) primary MEFs. Data are representative of three independent experiments. (B) Birth rates of OTULIN^+/+^, OTULIN^+/-^ and OTULIN^-/-^ offspring mice from OTULIN^+/-^ x OTULIN^+/-^ breeding couples. (C) Targeting scheme showing the LoxP-flanked (floxed) and *Otulin* liver parenchymal cell (LPC)-specific KO allele generated through Alfp-Cre-mediated recombination. The boxes indicate exons. LoxP sites are indicated by arrowheads. (D) Genotyping PCR showing different PCR fragments for each *Otulin* genotype (+, WT; FL, floxed).

**Supplementary Figure 2.** (A) Liver weight to body weight ratio in 10 week-old control (WT, n = 11) and OTULIN^LPC-KO^ mice (n = 8). Data are presented as mean ± SEM. **, p < 0.01. (B) Macroscopic appearance of 10 week-old OTULIN^LPC-KO^ and wild-type (WT) livers.

**Supplementary Figure 3.** (A) Lymphocyte infiltration and oval cell hyperplasia (red arrowheads) in liver sections from 10 week-old OTULIN^LPC-KO^ mice. 20x magnification. (B) Hepatocellular hypertrophy and irregular trabecular pattern. 20x magnification. (C) Mitotic figures (black arrows) and apoptotic cells (red arrowheads) in liver sections from OTULIN^LPC-KO^ mice. 40x magnification. (D) Cytoplasmic hepatocellular vacuolization in liver sections from OTULIN^LPC-KO^ mice. 20x magnification.

**Supplementary Figure 4.** (A,B) Representative flow cytometry plots and indicated gating strategy for live CD45+ cells isolated from livers of control (WT, A) and OTULIN^LPC-KO^ (B) mice aged 30 weeks. (C) % of monocytes, neutrophils, total macrophages and total cDCs as a % of live CD45+ cells as indicated. (D) % of each indicated macrophage population as % of total macrophages. ***, p < 0.005; ****, p < 0.001. Data are from 1 experiment with n = 4-5 per group.

**Supplementary Figure 5.** (A-B) Cytokeratin 19 (CK19) (A) and Sirius Red (B) quantification on liver sections from 10 week-old wild-type (WT, n = 5) and OTULIN^LPC-KO^ mice (n = 5) demonstrating oval cell hyperplasia and fibrosis in OTULIN^LPC-KO^ livers. CK19 and Sirius red stains evaluated by semi-quantitative grading (0, none/minimal; 1, mild; 2, moderate; 3, severe). (C) Relative mRNA level of *Tgfb1* in total liver lysates from 10 week-old OTULIN^LPC-KO^ mice (n = 6) and control (WT, n = 7) mice. Data are presented as mean ± SEM. *, p < 0.05.

**Supplementary Figure 6.** (A) Macroscopic appearance of 30 week-old OTULIN^LPC-KO^ livers. (B) Relative mRNA levels of HCC marker genes alpha-fetoprotein (AFP) and connective tissue growth factor (CTGF) in total liver lysates isolated from 10 week-old wild-type (WT, n = 11) and OTULIN^LPC-KO^ (n = 12) mice. (C) Relative mRNA levels of hepatic stem cell marker genes glypican 3 (GPC-3) and CD133 in total liver lysates isolated from 10 week-old wild-type (WT, n = 11) and OTULIN^LPC-KO^ (n = 12) mice. Data are presented as mean ± SEM. *, p < 0.05; **, p < 0.01. (D) Macroscopic appearance of 1 year-old control (WT) and OTULIN^LPC-KO^ liver. Scale bar, 5 mm. (E) Serum ALT, AST and ALP levels in control (WT, n = 6) and OTULIN^LPC-KO^ (n = 17), mice. Data are presented as mean ± SEM. **, p < 0.01; ***, p < 0.001. (F) Liver weight to body weight ratio in 1 year-old control (WT, n = 6) and OTULIN^LPC-KO^ mice (n = 9). Data are presented as mean ± SEM. ***, p < 0.001. (G) Representative hematoxylin/eosin-stained liver section showing HCC in 1 year-old OTULIN^LPC-KO^ mice. 10 x magnification. T, tumor.

**Supplementary Figure 7.** (A) Representative images of liver sections from 10 week-old OTULIN^LPC-KO^ mice and control (WT) mice after TUNEL staining. 20x magnification. (B) Quantification of the number of TUNEL positive cells. Data are presented as mean ± SEM, n = 5 mice per genotype. * p < 0.05.

**Supplementary Figure 8. TNF, TNF-R1, MLKL or MyD88 deficiency do not prevent liver pathology in OTULIN^LPC-KO^ mice.** (A) Macroscopic pictures of representative livers from a 10 week-old OTULIN^LPC-KO^, OTULIN^LPC-KO^ TNF^KO^, OTULIN^LPC-KO^ TNFR1^KO^, OTULIN/MLKL^LPC-KO^ and OTULIN^LPC-KO^ MyD88^KO^ mouse. Scale bar, 2 mm. (B) Serum ALT, AST and ALP levels in control (WT, n = 49), OTULIN^LPC-KO^ (n = 22), OTULIN^LPC-KO^ TNF^KO^ (n = 6), OTULIN^LPC-KO^ TNFR1^KO^ (n = 14), OTULIN/MLKL^LPC-KO^ (n = 2) and OTULIN^LPC-KO^ MyD88^KO^ (n = 6) mice. Data are presented as mean ± SEM. *, p < 0.05; **, p < 0.01; ***, p < 0.001; ****, p < 0.0001. (C) Representative hematoxylin/eosin-, cleaved caspase 3- and Ki67-stained liver section from 10 week-old OTULIN^LPC-KO^, OTULIN^LPC-KO^ TNF^KO^, OTULIN^LPC-KO^ TNFR1^KO^, OTULIN/MLKL^LPC-KO^ and OTULIN^LPC-KO^ MyD88^KO^ mice. Scale bar H&E, 100 μm, cleaved caspase 3 and Ki67, 200 μm. (D) Quantification of the number of cleaved caspase-3 positive cells in liver sections from control (WT, n = 7), OTULIN^LPC-KO^ (n = 6), OTULIN^LPC-KO^ TNF^KO^ (n = 4), OTULIN^LPC-KO^ TNFR1^KO^ (n = 8), OTULIN/MLKL^LPC-KO^ (n = 2) and OTULIN^LPC-KO^ MyD88^KO^ (n = 5) mice. Data are presented as mean ± SEM, *, p < 0.05, ** p<0.01, ***, p< 0.001. (E) Quantification of the number of Ki67 positive cells in liver sections from control (WT, n = 9), OTULIN^LPC-KO^ (n = 6), OTULIN^LPC-KO^ TNF^KO^ (n = 5), OTULIN^LPC-KO^ TNFR1^KO^ (n = 7), OTULIN/MLKL^LPC-KO^ (n = 2) and OTULIN^LPC-KO^ MyD88^KO^ (n = 6) mice. Data are presented as mean ± SEM. *, p < 0.05; ** p < 0.01, **** p < 0.0001.

