## Supplementary material for "OTULIN prevents liver inflammation and hepatocellular carcinoma by inhibiting FADD- and RIPK1 kinase-mediated hepatocyte apoptosis": Supl. Figures

A.

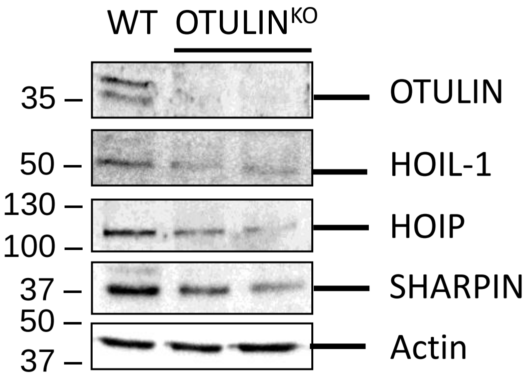


B.

| OTULIN$^{+/-}$ x OTULIN$^{+/-}$ | Expected | Observed (at birth) |
|---|---|---|
| OTULIN$^{+/+}$ | 25 % (9-10) | 34 % (13) |
| OTULIN$^{+/-}$ | 50 % (19) | 66 % (25) |
| OTULIN$^{-/-}$ | 25 % (9-10) | 0 % (0) |
| Total (observed) | 100 % (38) | 100 % (38) |

C.

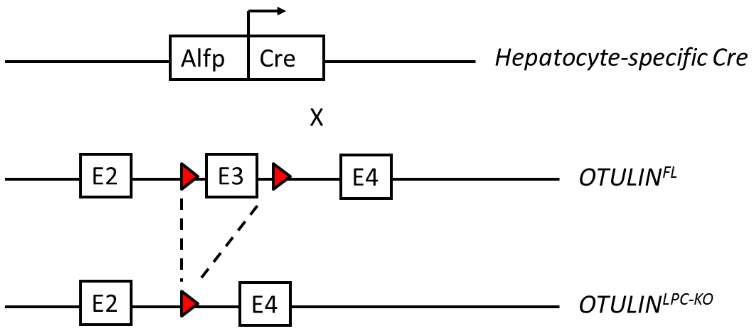


D.

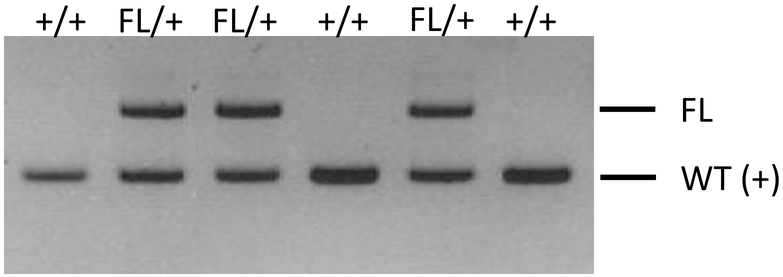


Supplementary Figure 1

A.

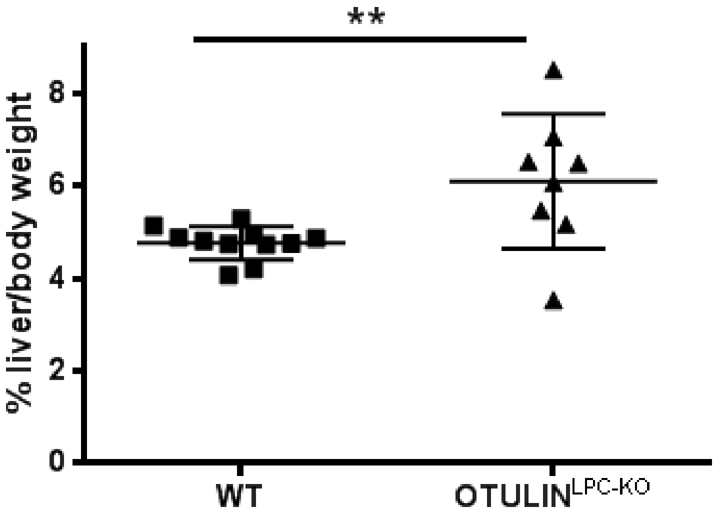


B.

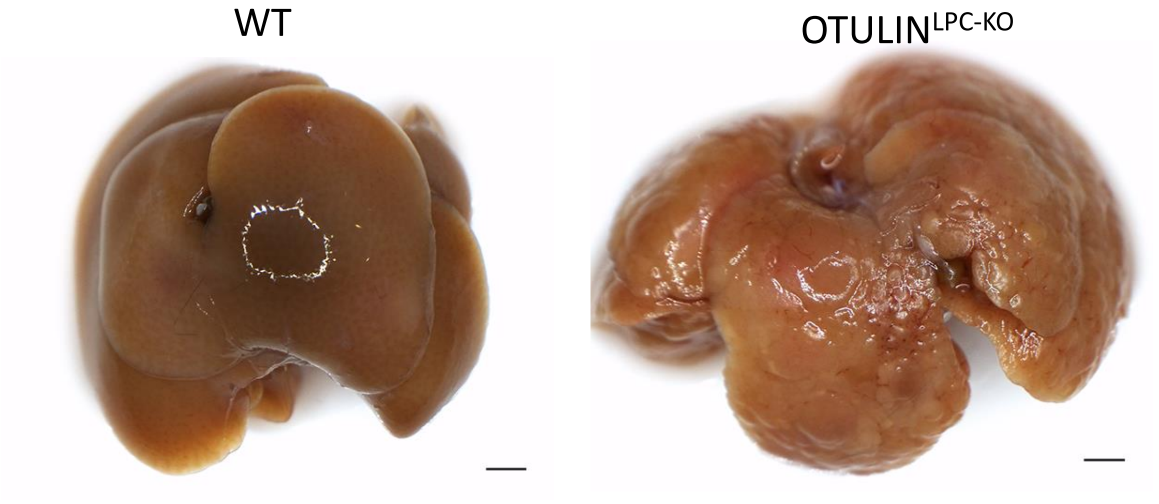


Supplementary Figure 2

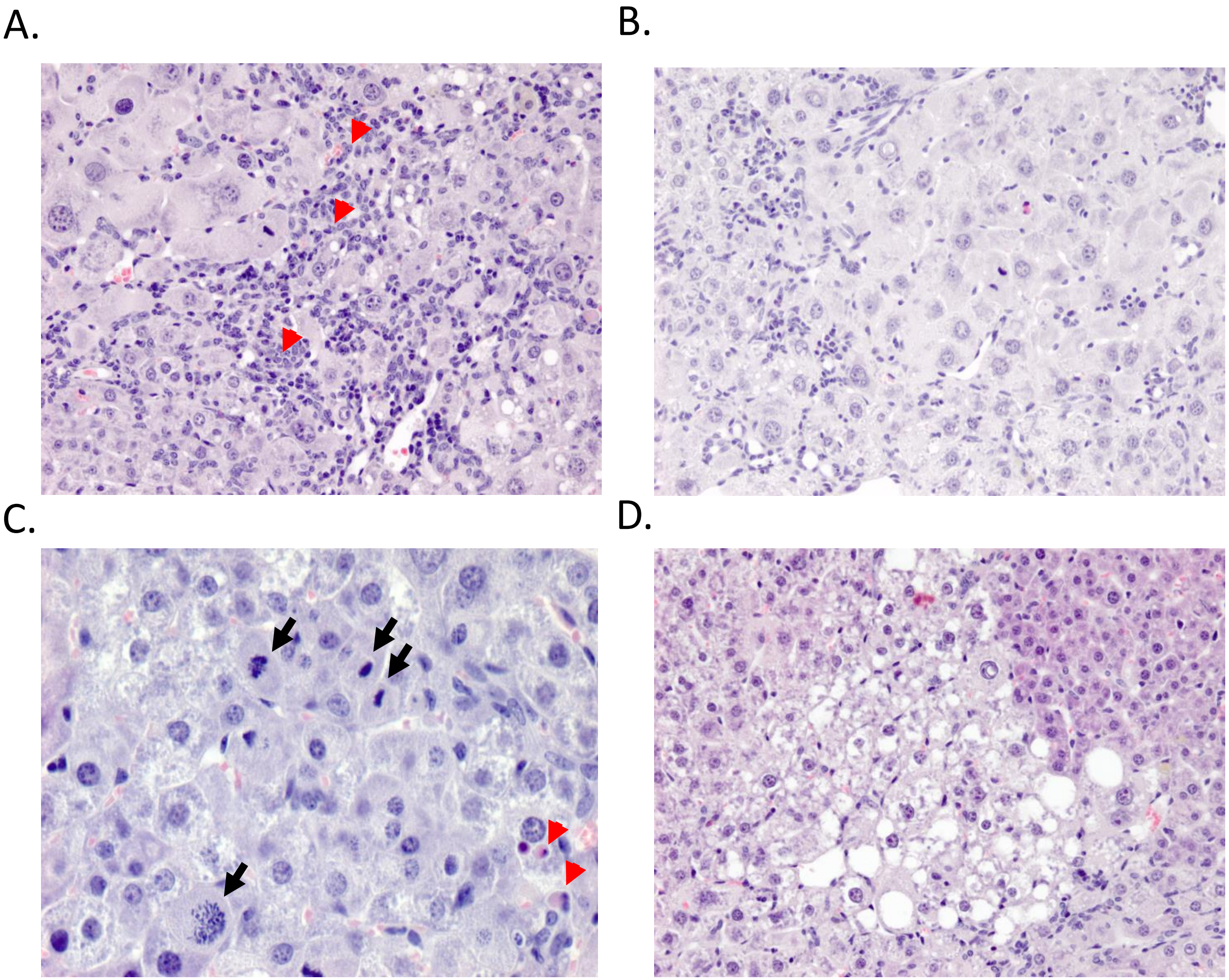


Supplementary Figure 3

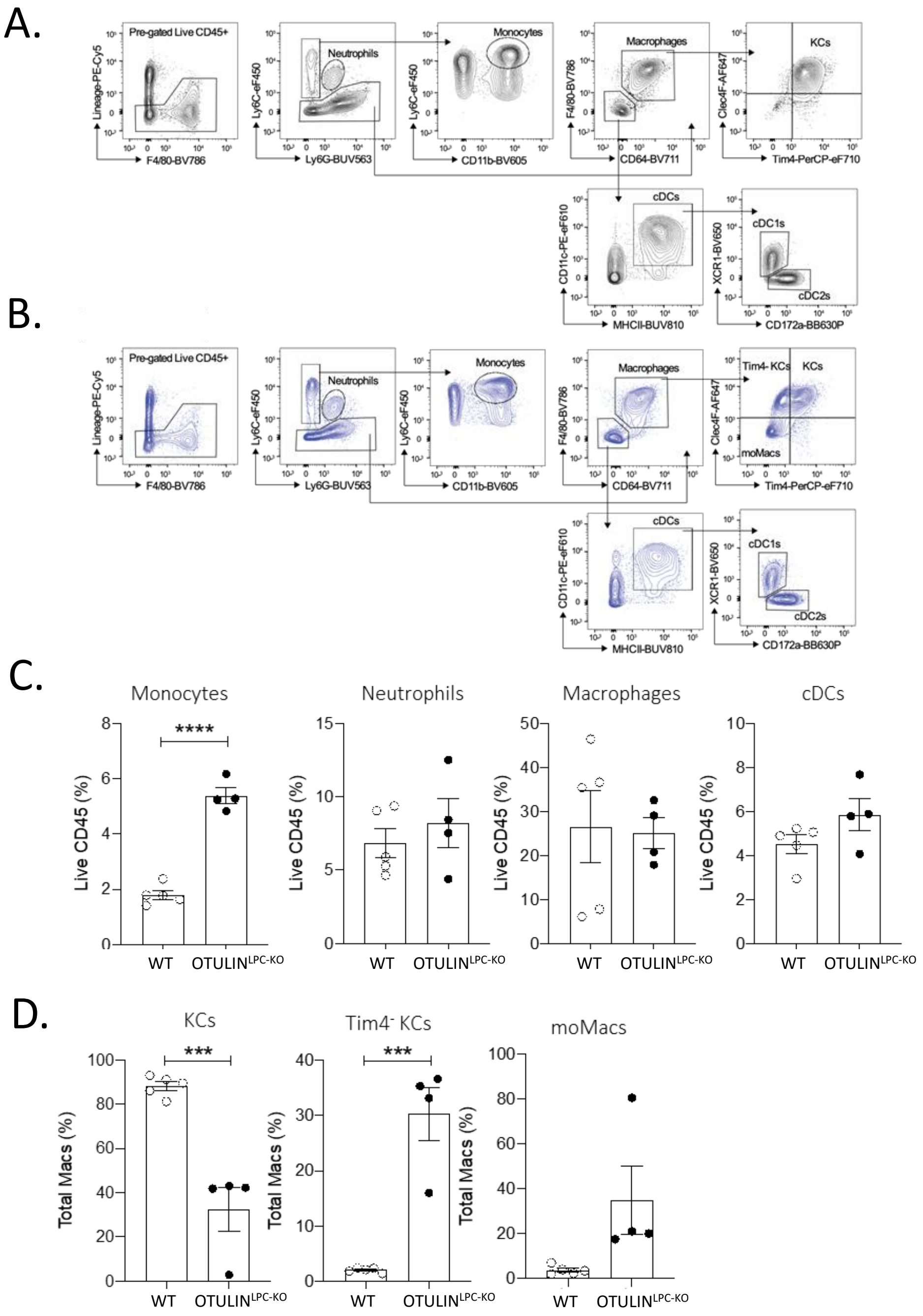


Supplementary Figure 4

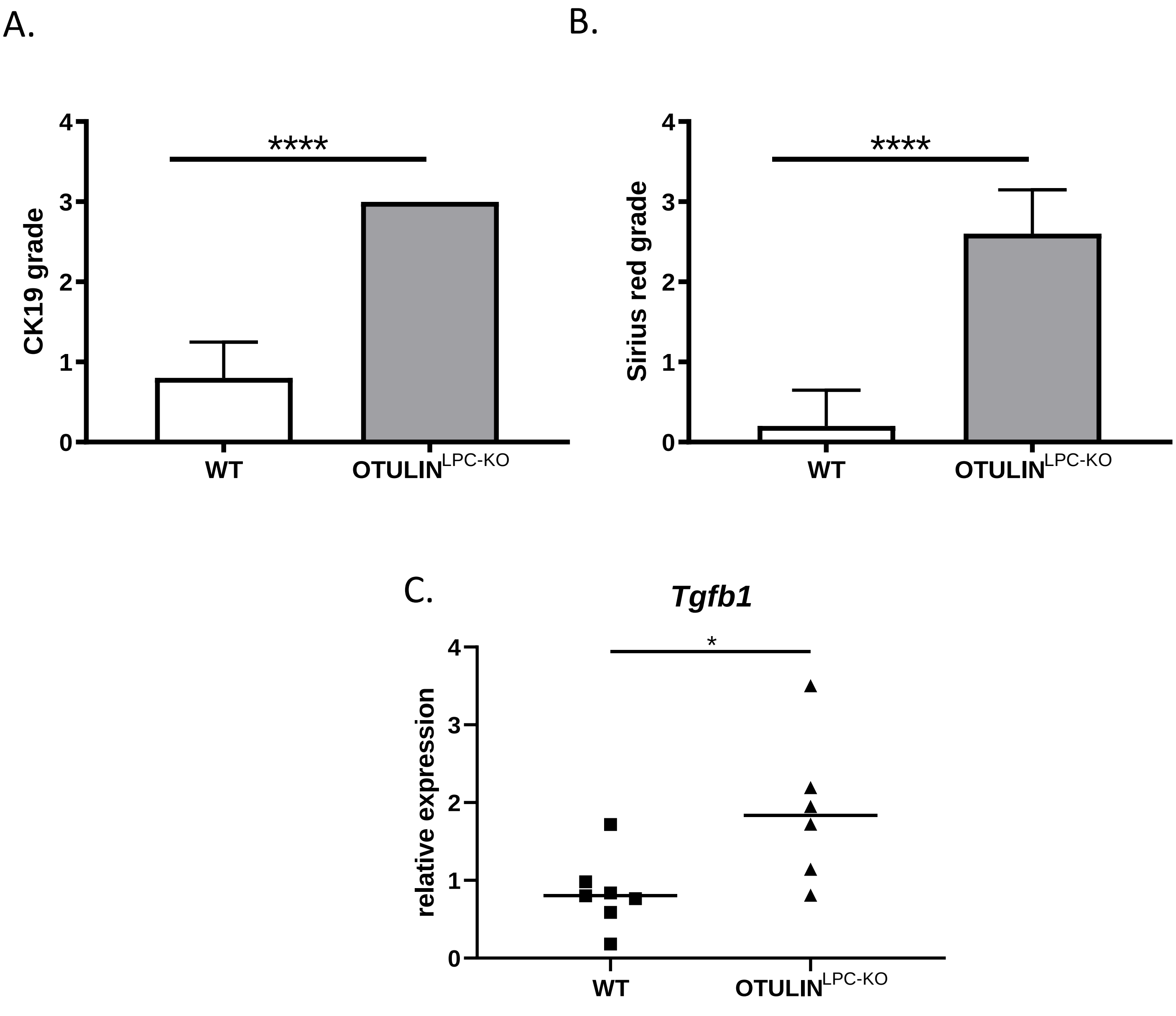


Supplementary Figure 5

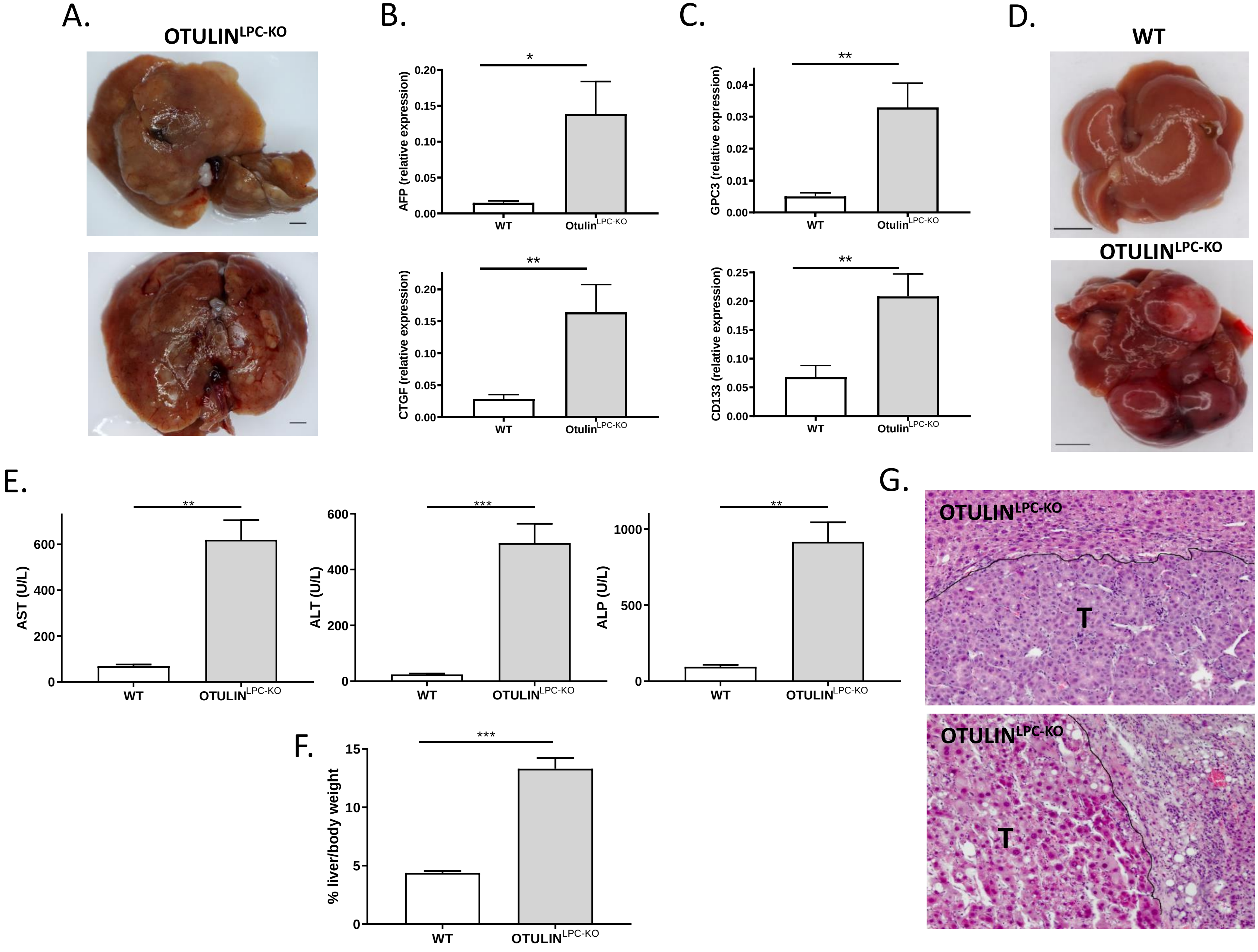


Supplementary Figure 6

A.

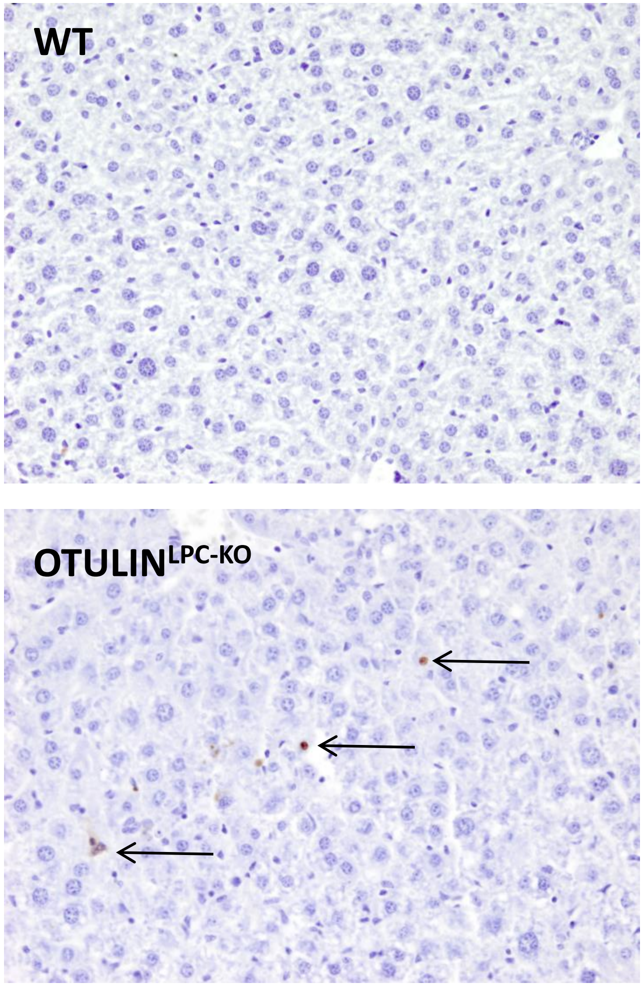


B.

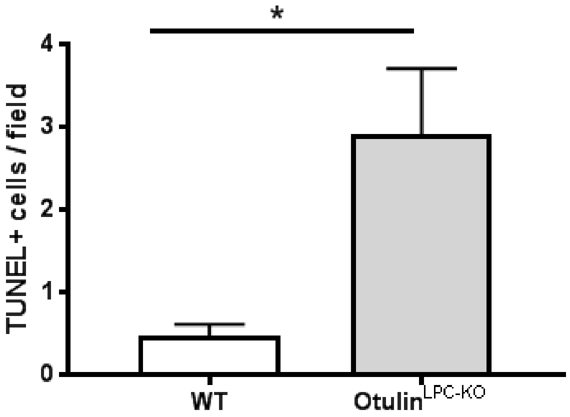


Supplementary Figure 7

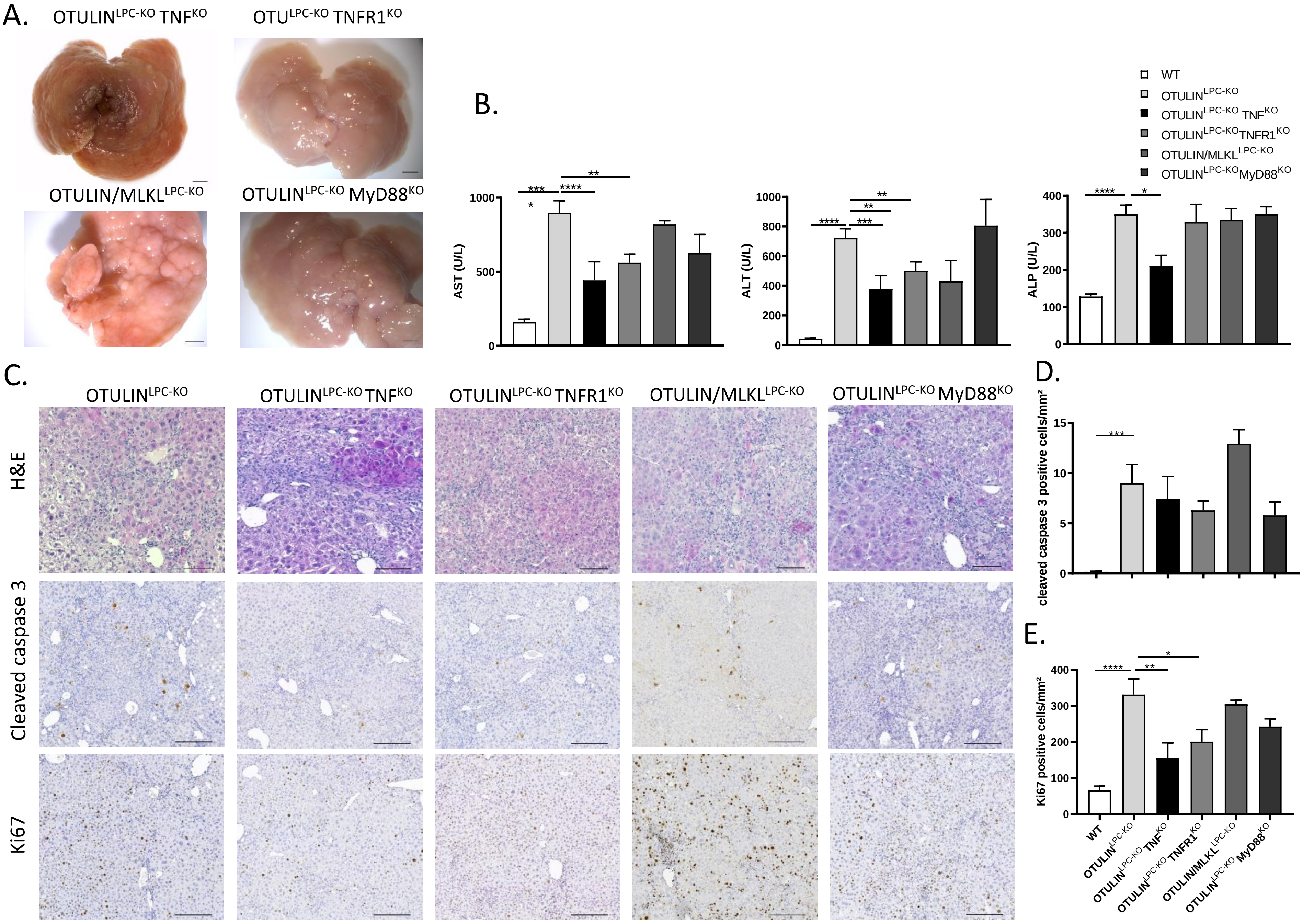


Supplementary Figure 8
